# Computational framework for statistical epistasis supports XOR penetrance function in a living system

**DOI:** 10.1101/2023.05.09.539940

**Authors:** Sandra Batista, Vered Senderovich Madar, Philip J. Freda, Priyanka Bhandary, Attri Ghosh, Apurva S. Chitre, Abraham A. Palmer, Jason H. Moore

**Author notes:** These authors contributed equally to this work (Sandra Batista, Vered Senderovich Madar, Philip J. Freda). These authors also contributed equally to this work (Abraham A. Palmer, Jason H. Moore).

## Abstract

Statistical epistasis has been studied extensively because of its potential to provide evidence for genetic interactions for phenotypes, but there have been methodological limitations to its exhaustive, widespread application. We present new algorithms for the interaction coefficients for standard regression models for epistasis that permit many varied encodings for the interaction terms for loci and efficient memory usage. The algorithms are given for two-way and three-way epistasis and may be generalized to higher order epistasis. Statistical tests for the interaction coefficients are also provided. We also present an efficient matrix based algorithm for permutation testing for two-way epistasis. We offer a proof and experimental evidence that methods that look for epistasis only at loci that have main effects may not be justified. Given the computational efficiency of the algorithm, we applied the method to a rat data set and mouse data set, with at least 10000 loci and 1000 samples each, using the standard Cartesian encoding and the XOR penetrance function for the interactions, to test for evidence of statistical epistasis for the phenotype of body mass index. This study revealed that the XOR penetrance function found greater evidence for statistical epistasis in many more pairs of loci in both data sets and in the rat data set, those pairs of loci found using the XOR penetrance function are enriched for biologically relevant pathways.

**Author summary:** Epistasis, the interaction between two or more genes, is likely integral to the study of genetics and present throughout nature. Yet, it is seldom fully explored as most approaches primarily focus on single-locus effects (such as GWAS), partly because analyzing all pairwise and higher-order interactions requires significant computational resources. Many current methods for epistasis detection only consider a Cartesian encoding for interaction terms. This is likely limiting as epistatic interactions can evolve to produce varied relationships between genes, some non-linear. In this work we describe computationally efficient algorithms for the detection of statistical epistasis that allow for varied interaction encodings for modeling epistasis. Our methodology efficiently detects pairwise and three-way epistatic interactions in two closely related species (rat and mouse) under both Cartesian and XOR interaction encodings. Our results in both species show that many biologically relevant epistatic relationships would have been undetected if only one interaction encoding was applied providing evidence that more varied models for interaction may need to be applied to describe epistasis that occurs in living systems.

## Introduction

Epistasis is challenging to detect yet likely widespread and integral in biology. Evidence for epistasis has been discovered in a host of biological systems and phenotypes including mandible size in mice [1], cardiovascular disease susceptibility [2], coronary artery restenosis [3], cystic fibrosis [4, 5], and sporadic breast cancer [6] in humans, and most recently and robustly in two studies investigating non-additive genetic effects in yeast [7, 8]. The yeast studies have collectively identified thousands of epistatic two-way and three-way interactions that vary across lineages and growth conditions. Additionally, they have identified large epistatic hubs that are involved in most interactions detected In the most recent study in yeast, non-additive effects accounted for one-third of the broad-sense heritability [8]. These studies provide strong evidence that epistasis accounts for a large portion of the non-additive genetic variation in natural systems.

Since loci may contribute to phenotypes through nonlinear interactions, epistasis may account for genetic variation not explained by single-locus approaches. Indeed, it has been shown that the main effect of one SNP can significantly change when the allele frequencies in a second SNP are altered [9]. Examples of where epistasis may have a crucial role include biomolecular interactions in gene regulation, signal transduction, and biochemical networks [10–12]. Thus, many phenotypes can be viewed as the result of vast interconnected biological networks and systems [13–15]. These biological systems likely arise to form compensatory networks that aid in buffering against genetic and environmental change (i.e., canalization) [16–19]. It is likely that, at the core of these networks, interactions among polymorphisms from multiple pathways exist and are integral to organismal development, homeostasis, and survival [17].

Robust methodologies aimed at detecting and describing statistical epistasis are required to investigate genotype-phenotype associations and disease susceptibility [11]. Since experiments that could biologically validate detected statistical epistatic interactions are rare [13–15], the development of these methodologies will assist in initiating further scientific exploration. In this work, we examine how to efficiently compute linear regression models for epistasis that permit varied encodings of the interactions of loci and provide statistical evidence for epistasis. Recent computational and theoretical work has presented a new way to calculate each of the coefficients of a linear regression model [20]. In this work we aim to demonstrate the usefulness and practical significance of the closed forms in general and more specifically in genetic studies and in particular, epistasis. We present algorithms for providing statistical evidence for two-way and three-way epistasis using standard models for epistasis using these closed forms. These algorithms may be used efficiently on subsets of loci or genome-wide, are entirely parallelizable, provide statistical tests, and permit flexibility in encoding the interactions of loci. Many methods for interaction terms only consider the Cartesian encoding that essentially multiplies the genotypes at two loci together. In our method any function may be applied to interacting genotypes and we demonstrate this using the exclusive or (XOR) encoding penetrance function. We also discuss how these algorithms may be generalized for higher order interactions. Permitting many types of encodings for interactions allows for nonlinearity to be included in the traditional statistical models for epistasis and for biologists to use a variety of penetrance functions to model epistasis.

As specific examples, we apply our algorithms to detect statistical evidence for two-way and three-way interactions using the Cartesian and XOR penetrance functions on the phenotype of body mass index (BMI) in real data sets from rats (*Rattus norvegicus*) from a GWAS study [21–23] investigating obesity-related traits and from mice (*Mus musculus*) from Wellcome Trust [24, 25]. For both data sets we use approximately 10,000 SNPs for two-way epistasis detection using both penetrance functions. The implementations of these algorithms given in Python are offered via GitHub^1^.

### Computational Challenges in Detecting Epistasis

Methods to explore genotype-phenotype associations and detect epistatic interactions are often computationally intensive and may completely ignore non-additive effects. Single-locus analyses like genome-wide associations studies (GWAS) can detect strong main effects, but face difficulties when applied to combinations of variables for many reasons [14, 15]. The first is that multi-locus genotype (MLG) combinations have smaller representative samples compared to the original data set. Second, in most approaches that attempt to model interactions using linear models, interactions are only considered when significant main effects of variables are identified [14, 15]. Although it is tempting to expect loci with significant main effects to also be involved in interactions, there is no statistical justification for this. Third, linear models have greater power to detect main effects compared to interactions [26–28]. Fourth, many linear model approaches model interactions using the Cartesian product for ease of computation whereas other models for interactions may also be plausible. We expand more on this in the following section. Finally, when considering higher order interactions, as the number of loci in k-wise combinations increases, the number of variables in the standard regression models increases exponentially and the total number of sets of loci of size *k* to consider of all *n* possible loci increases polynomially as more loci are considered. This exhaustive search space creates issues with computational tractability as investigating pairwise and higher-order interactions becomes extremely difficult to achieve efficiently. Given these challenges, some of which are inherent in exhaustively considering all possible subsets for k-wise interactions in a set of n loci, many methods apply various techniques to reduce this search space or to exploit parallelism in underlying matrix libraries for computational efficiency.

There are many techniques to reduce the search space for epistasis algorithms. One example, the multifactor dimensionality reduction (MDR) technique finds multilocus genotypes that have high or low association with disease and defines new variables that explain the relationship of both loci [6, 29, 30]. MDR can be combined with other machine learning methods and has been extended to handle population structure [31]. In an approach that bypasses the conventional search space of epistasis algorithms, Crawford et al. developed the “MArginal ePIstasis Test” (MAPIT) method that measures the marginal epistatic effect of a single loci against all other loci all at once [32]. MAPIT can be used as a way to screen for all loci that have significant marginal epistastic effects for subsequent tests to see which pairs of loci may be involved in epistasis. More comprehensive recent surveys on epistasis are offered by Ogbunugafor and Scarpino on higher order epistasis [33], Niel et al. on statistical and computational challenges of varied approaches [34], and Russ et al. for performance comparison of many varied epistasis detection methods [35].

In exhaustive methods for statistical epistasis for discrete (case/control) phenotypes, a common approach to make the problem more computationally tractable is to use contingency tables for approximating pairwise epistasis. One of the most commonly used software packages for applying the standard pairwise epistasis regression models is PLINK [36]. In PLINK pairs of loci are scanned using an approximate method that evaluates the Z-scores for the odds ratio at the loci between cases and controls or BOOST that uses bitwise operations for calculating the likelihood ratio test [37]. It is then possible to apply the entire regression to the pairs of loci that pass the screening stage. Zhang et al. use minimum spanning trees to update contingency tables for test statistics for evaluating pairs of loci for epistasis in the TEAM algorithm [38]. These aforementioned approaches work for pairwise epistasis. Bayat et al. offer the BitEpi method for handling up to four-way epistasis using bit efficient counting for contingency tables, an entropy metric to calculate the interaction effect, and permutations for p-value calculations [39].

Another approach to ease the computational burden for exhaustive methods for statistical epistasis is to exploit the parallelism from hardware architectures or underlying matrix operations. Schupbach et al. present FastEpistasis as an extension to PLINK that for quantitative phenotypes calculates the interaction term for a pair of loci by using the QR decomposition to solve the ordinary least squares [40]. The efficiency from their method is from the parallelism of the calculations and the hardware architectures. Zhu and Fang offer the method MatrixEpistasis that evaluates the standard regression model for epistasis for the interaction term by computing the residuals of the linear regression model between the phenotype and loci and the residuals for the regression model between the interaction and loci [41]. Like our method some of the efficiency of MatrixEpistasis is achieved by only focusing on the interaction term and corresponding test statistic, but the efficiency of their method also relies on the efficiency of underlying matrix operations and accordingly the method can only handle Cartesian encodings of the interaction term. The work of Zhu and Fang also considered covariate adjustment, but that may be done before applying our algorithm as is essentially done in their implementation and commonly done in practice as well.

Unlike these other methods, our method permits many different encodings for the interaction, thus potentially introducing some nonlinearity into the model. Our method also explicitly deals with higher order epistasis. The method we present is the result of new computational and algorithmic insight for solving the ordinary least squares that expresses the estimates in closed forms. As a result our method is also very efficient in terms of memory and entirely parallelizable for testing each subset of loci for epistasis.

### The XOR Penetrance Function

It is not fully understood how epistatic interactions evolve, are maintained, or are structured in biological systems considering that methodologies specifically designed to systematically identify non-linear interactions are not common [13–15]. As an example of the possible complexity of epistasis, Li and Reich propose over a hundred full penetrance interaction models, some of which are not linearly separable [42]. The possible types of interactions become infinite when we consider incomplete penetrance. To test our methodology’s capability of supporting multiple interaction encodings when detecting epistasis, we applied the exclusive or (XOR) penetrance function encoding in addition to Cartesian encoding (File S2). The XOR function is described by Li and Reich as model M170 [42]. We chose XOR because of its extreme difference compared to a standard Cartesian encoding. Under the full penetrance XOR model, the phenotype is entirely dependent on the multi-locus genotype. Thus, in it’s purest form, XOR is not linearly separable or detectable using any single-locus analyses like GWAS. Due to these aspects, the XOR model is commonly considered to not be biologically plausible. Despite being an extreme example of epistasis, models that could exist in a biological system and are consistent with the XOR model have been postulated [43, 44]. Detection of statistically significant epistatic interactions that are consistent with the XOR model in a biological system may justify investigating more complex genetic architectures in nature. Furthermore, detection of statistically significant epistatic interactions using the XOR model may provide justification to design validation experiments to identify biological XOR epistasis in living systems. We demonstrate that the XOR encoding detects statistical epistatic interactions not identifiable by a Cartesian model.

## Methods

### Models for Epistasis

At the center of modeling epistasis is regression and we may now consider whether locus *i* and locus *j* have an epistatic interaction, *I*_*i,j*_, affecting the phenotype, *p*, by considering their interaction effect term, *β*_3_ :

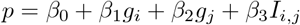

We may remove the intercept and simplify the computation of the model by subtracting off the means of the variables, i.e. 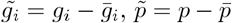 and 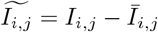 :

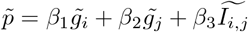

We are only concerned with *β*_3_, the coefficient for the interaction term, 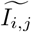, in the model because it represents the partial correlation between the phenotype and the interaction term given both loci. It is important to note that there is flexibility in this model in how the interaction is encoded. Although it is common to multiply values of the loci for each sample as is commonly done for the Cartesian encoding for ease of computation, many different encodings may be used as we demonstrate for the XOR penetrance function (File S2). While the Fisher t-statistics for hypothesis testing can be applied (e.g. [41]), a direct *T* test for a regression coefficient may also be applied instead

The model for epistasis may be generalized to higher order interactions. For *k*−order interactions between a set of *k* loci, *g*_*i*_ for 1≤ *i* ≤*k*, we must consider the interactions of all subsets of the loci from the empty set to the entire set of all *k* in a manner reminiscent of applying the binomial theorem where the coefficient is the interaction rather than binomial coefficient. For the epistatic interaction of a subset, we will use the variable *I*_*s*_ where *s* is the subset. An appropriate function, *f*, for the encoding, such as XOR or Cartesian, may be applied to the sets as well as the interaction term of the entire set of *k* loci. All variables may then also be centered by subtracting off the means The model for the epistatic interactions affecting phenotype *p* may be expressed as follows:

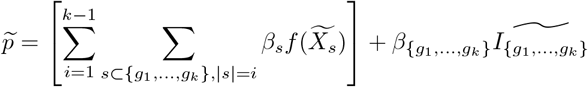

The model that we will consider for 3-way epistasis for loci *g*_1_, *g*_2_, *g*_3_ is accordingly after centering the variables:

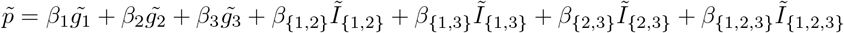

For detecting epistasis the interaction term of interest is *β*_{1,2,3}_ for 3-way epistasis and 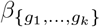 for k-way epistasis. It is important to note that for *k*−way epistasis the regression must include *O*(2^*k*^) variables for each set of *k* loci and that to check for all possible *k* −wise interactions between *n* loci, there are *O*(*n*^*k*^) sets that must be checked for the given model. For a given set of *k* loci, we explore how it is not necessary to find the interaction coefficients for all 2^*k*^− 1 variables and may focus only on the final coefficient of interaction along with more efficient and powerful statistical tests of interaction. We give algorithms explicitly for *k* = 2, 3 and consider how these may be generalized.

### Estimation and Statistical tests for interaction

The conditional association between the phenotype, *p*, and the interaction term, *I*, while conditioning upon the main effects of the loci, **Z** = {*g*_*i*_, *g*_*j*_ }, is the epistasis interaction. This may be computed in multiple ways. The first way is to compute the partial correlation coefficient between *p* and *I*, as the dot product between two standardized residuals: the residuals for the regression model between the phenotype and the loci and the residuals for the regression model between the interaction and loci This approach requires the computation of two vectors of residuals estimated by two linear regression models. The first linear regression model is between the phenotype and loci, *p* ∼*β*_0_ + **Z** and the second model is between the interaction and loci, *I* ∼*β*_0_ + **Z**, as was done in the work of Zhu and Zhang [41]. The standardized residual of the linear regression model between the phenotype and loci, *p* ∼ *β*_0_ + **Z**, is 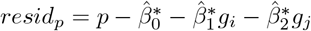 where the 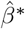 are the estimates from the model. The standardized residual of the regression between the interaction and loci, *I* ∼ *β*_0_ + **Z**, is 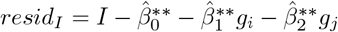 where again the 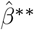 are the estimates from the model. This means two separate regression models are applied as was done in the work of Zhu and Zhang [41]. The partial correlation coefficient between *p* and *I* is thus *r* :

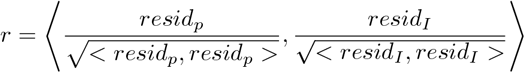

The second way is to compute a single regression coefficient, the corresponding regression coefficient, 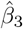 in the model between the phenotype, loci, and interaction term: *p*∼ *β*_0_ + *β*_2_*g*_1_ + *β*_2_*g*_2_ + *β*_3_*I*. Both measures are variants of each other, as a computed *r* = 0 will necessarily imply a computed 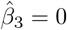 and vice versa. This is due to Yule’s equivalence formula [45],

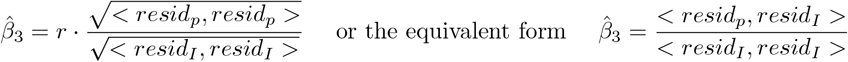

We know now that *< resid*_*p*_, *resid*_*I*_ *>*=*<* **p**, *resid*_*I*_ *>* and know a direct way to estimate 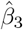 and each of the other coefficients that is the crux of our algorithm [20]. Consider the estimation of the model assuming all variables are mean centered:

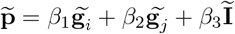

We have by considering the residuals of each loci with the interaction term an expression for the interaction effect term, *β*_3_:

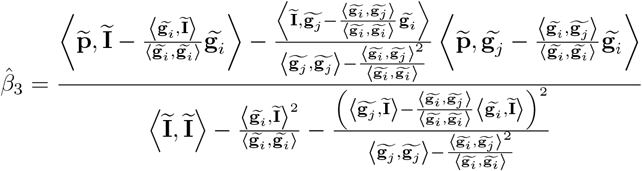

In particular, we may write the vector of partial residuals of the interaction term with each loci, *resid*_*I*_, directly as

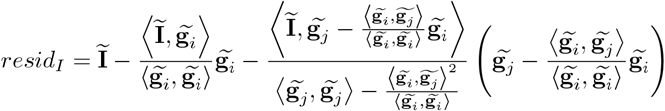

while in a similar way we may write the vector of partial residuals of the phenotype with each loci, *resid*_*p*_, directly as

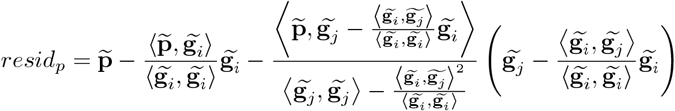

As a result we may rewrite the interaction effect term, 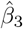. It may be expressed in terms of the ratio of two dot products between the dependent variable and the partial residuals and between the independent variable *I* and partial residuals as follows:

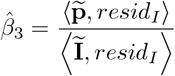

As for hypotheses test, we test the null hypotheses

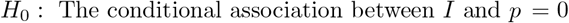

against the alternative

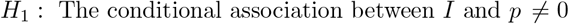

This calls for multiple testing correction to protect against the inflated Type I error and we use the Benjamini Hochberg false discovery rate (FDR) controlling procedure for this. The statistical test will be a T test with *n*− *v* −1 as degrees of freedom where *n* is the number of samples and *v* is the number of parameters in the model.

Zhu and Zhang address the partial correlation between the interaction term *I* and the phenotype *p*, and compute it using the dot product of the residuals [41]. As for a statistical test to test for null value for the association, the Fisher approximate test is then used [46, pp. 26]:

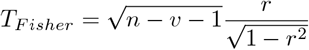

The T test statistic for testing the null hypothesis against the alternative will be in terms of successive residuals: the partial residual, *resid*_*p*_ aforementioned, and the global residual, 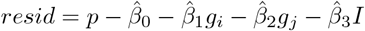, for *p* ∼*β*_0_ + *β*_2_*g*_1_ + *β*_2_*g*_2_ + *β*_3_*I* after adding *I* to the model. We observe that we may calculate the mean sum of squares or the mean square error, MSE, using the global residual, that is,

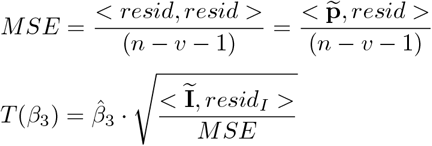

As a result we may calculate the test statistic as well:

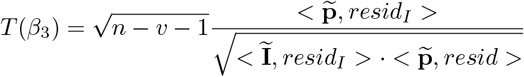

Both of the T tests above require the computation of the 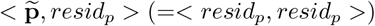 or the MSE (for modeling *p*) or where it is possible to verify that

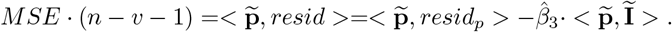

### Algorithms for Epistasis Detection

We may now give an algorithm for computing the interaction coefficient and test statistic for epistasis for two loci, *g*_*i*_ and *g*_*j*_. We may encode their interaction, *I*_*i,j*_ using any type of epistasis that may be suspected and often by default the Cartesian encoding (or product) is used. We will use the centered variables for our algorithm where the means have been subtracted, i.e. 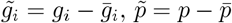 and 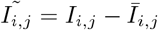. The algorithm residualizes the first locus from the second and then each loci from their interaction in order. The algorithm does the same in residualizing each loci from the phenotype in order. Finally to calculate the residuals of the entire model we residualize the interaction from the phenotype. In the case of pairwise epistasis, the algorithm only requires the centered variables to calculate several vector dot products and vector additions (subtractions). In the exposition given below we calculate the direct t-test statistic for the regression coefficient. Let the number of samples for each loci and the phenotype be *m*.

#### Algorithm 1

Interaction Coefficient for Pairwise Epistasis

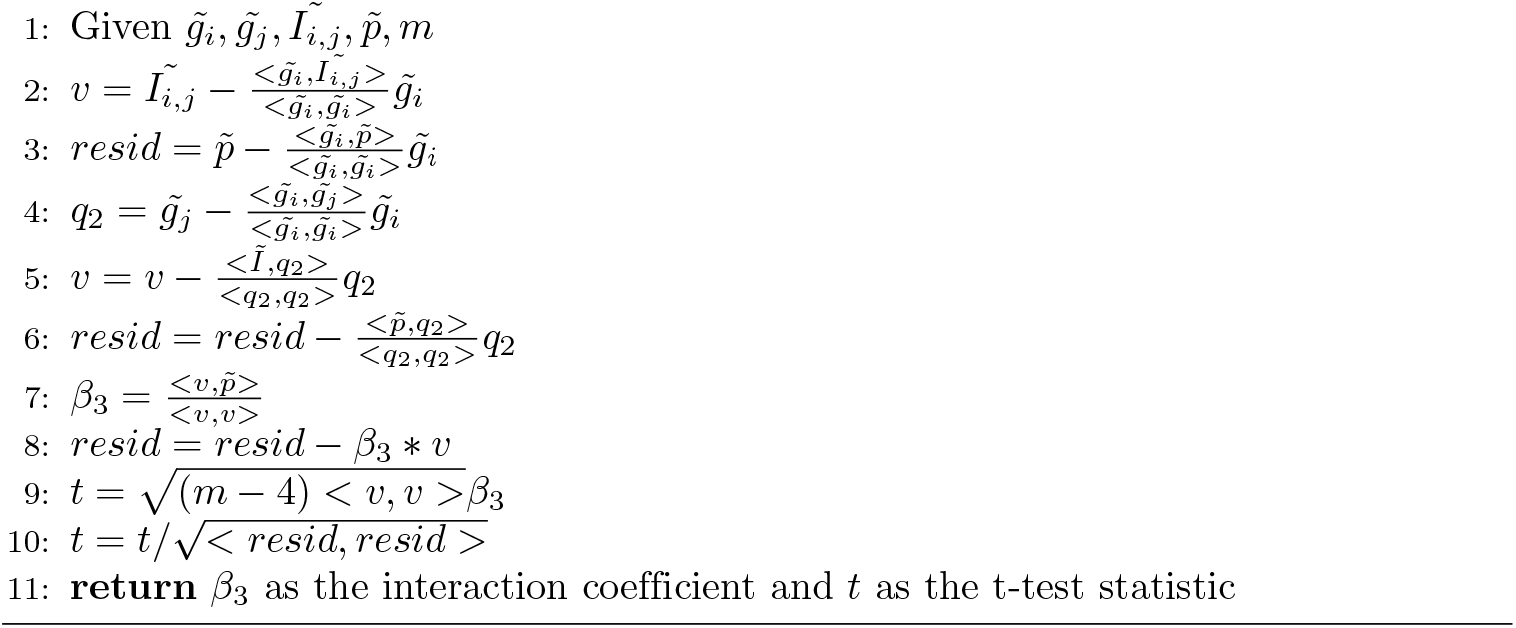

The algorithm is linear in the number of samples *m*. However, if the aim is to test all possible pairwise epistasis for *n* loci, this will still be an *O*(*n*^2^*m*) algorithm. The flexibility here is to test any pair of loci for epistasis with a variety of test statistics and a variety of possible interaction encodings. There is also the practical concern regarding floating point representation in implementations if any of the vector norms or norms squared, such as the norm of the residuals (in line 10), are close to zero. In some implementations this could raise an error because of division by zero, for example, because of limitations in floating point representations. In this case we recommended that any such errors that arise, while not errors in the algorithm or implementation themselves, but merely limitations of floating point representations, be logged for further inspection as we have done in our experiments.

We will now consider the algorithm for 3-way epistasis for a set of loci. While technically the algorithm is still linear in the number of samples since *k* = 3 is a constant, we begin to see more of the effects on the complexity of the number of variables in the model. For the case of 3-way epistasis, there are only 7 variables, but more generally there are *O*(2^*k*^) variables in models for *k* −way epistasis and the algorithm is quadratic in the number of variables while linear in the number of samples, so the more general case has complexity *O*(2^2*k*^*m*) for a single set of *k* loci. The models themselves introduce a number of variables exponential in *k* and for all possible sets of *k* loci from *n* the complexity is *O*(*n*^*k*^2^2*k*^*m*), so there is a need to prune the space of loci to check and variables in the models.

However, first we present the algorithm for 3-way epistasis detection and for the sake of exposition mention how it can be generalized to *k*− wise epistasis with the caveats we have given for its time complexity. For the encoding of all interactions we assume that a Cartesian encoding (or product of the loci) are used. We assume that we have three loci *g*_1_, *g*_2_, *g*_3_ and the phenotype, *p* as well as their encodings and they are all centered (with their means subtracted from them). First we construct an 8 by *m* matrix *Q* that will be used to residualize each variable against each other in the order given. For ease of exposition, we will index the columns of *Q* for each variable starting at 1 where 1 will be the index for 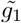, 7 for 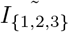 and 8 for 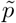. It is important to note that we are overwriting Q. This is important to mention since it clarifies not only the correctness of the algorithm, but highlights the space efficiency of these algorithms. Once the intermediate matrix for the 8 (or more generally 2^*k*^) variables for the model is constructed for which only read only access of the entire data matrix is needed, no additional access to the data matrix is needed and only additional constant space for several dot products and to hold the interaction coefficient and test statistic are needed

#### Algorithm 2

Interaction Coefficient for 3-way Epistasis

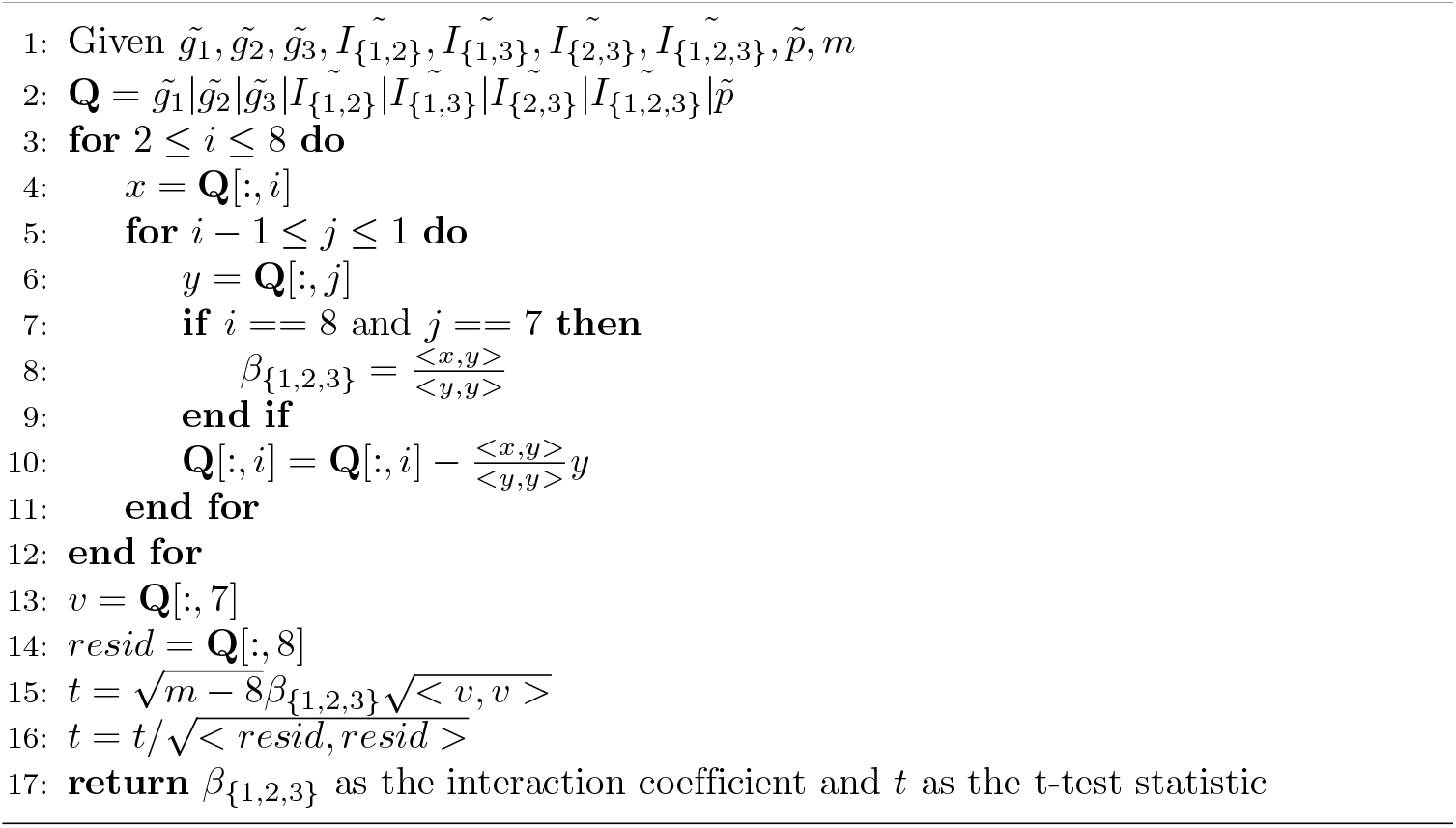

To generalize the 3-way epistasis algorithm to be a *k* wise epistasis algorithm, we need to first enumerate the 2^*k*^ variables of the model and center them by subtracting off their means to construct **Q**. Then it is necessary to residualize the variables by changing 8 to the value 2^*k*^ on line 3. To get the interaction coefficient for all *k* loci, *β*_{1,…,*k*}_, on line 7 check that *i* == 2^*k*^ and *j* = 2^*k*^− 1. To calculate the t-test statistic, on line 13 set *v* = **Q**[:, 2^*k*^− 1] and on line 14 *resid* = **Q**[:, 2^*k*^] and adjust the degrees of freedom on line 15. If we have a set of loci of interest, this algorithm and its generalization can calculate their interaction coefficient and test statistic efficiently. However, as we noted if we do not have a known set of loci a priori, then there is work to be done to prune what would be the exhaustive search space as we will consider in the next section.

The extension of the algorithms for permutation tests can be easily done by permutations of the dependent variable **p** while computing *resid*_*I*_, *resid*_*p*_ and in the case of a pairwise epistasis, 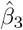 as

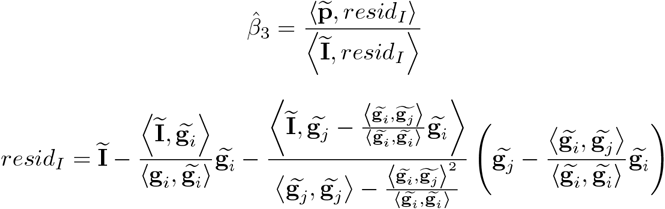

and

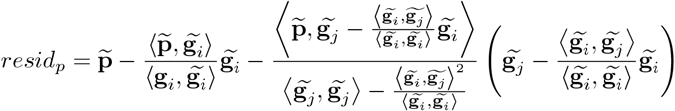

They may be used in order to obtain the test statistic

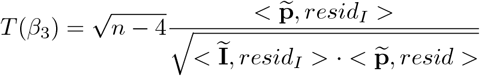

In the case of pairwise epistasis

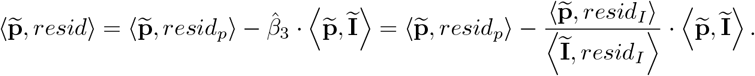

Note that *resid*_*p*_ and the dot product of the permuted phenotype and interaction term needs to be updated for each permutation. The interaction term and its residual may be reused for all permutations. The extension of the pairwise epistasis algorithm with permutation testing follows in Algorithm 3. The interaction coefficient, *β*_3_ and T statistic are computed as in Algorithm 1. Algorithm 3 calculates the interaction coefficient and test statistic for each permuted phenotype. The p-value is the percentage of permuted phenotypes with test statistics at least as large as the test statistic for the original phenotype. We present the algorithm using matrix calculations on the permuted phenotype matrix, *P*, an *m* by *K* matrix, where *m* is the number of samples and *K* is the number of permutations and each column of *P* is a permutation of the original phenotype vector, 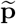. There are several advantages to this approach in that calculating *P* once for all permutation tests permits efficient parallelization of the permutation testing. The permuted phenotype matrix may be divided into submatrices of columns of permutations if need be especially for memory limitations. In this case the calculation of the p-value would be modified to be done across the divisions, i.e. return the number of tests at least as large original t-test in absolute value and calculate p-value after all computations are done. Second using matrix operations permits implementations of the algorithm to exploit underlying parallelization and efficiency of matrix libraries such as BLAS and LAPACK used by NumPy in Python.

It should be noted that permutation tests offer a robust alternative statistical test which is preferred in case there is suspicion that the required linear regression assumptions of nonlinearity or nonnormality fail to hold. Permutation tests are also known to control better the family wise error rate in the scope of the specific model under testing. As each permutation test is performed under a specific epistasis model, when considered altogether, for all pairs, the control of the global inflation of the Type I error is not assured. To determine if permutation tests notably affect our results compared to FDR correction alone, we implemented Algorithm 3 and applied it to perform 1000 permutations for pairwise epistasis detection using the XOR encoding on the rat data set.

#### Algorithm 3

Permutation Test P-value for Pairwise Epistasis

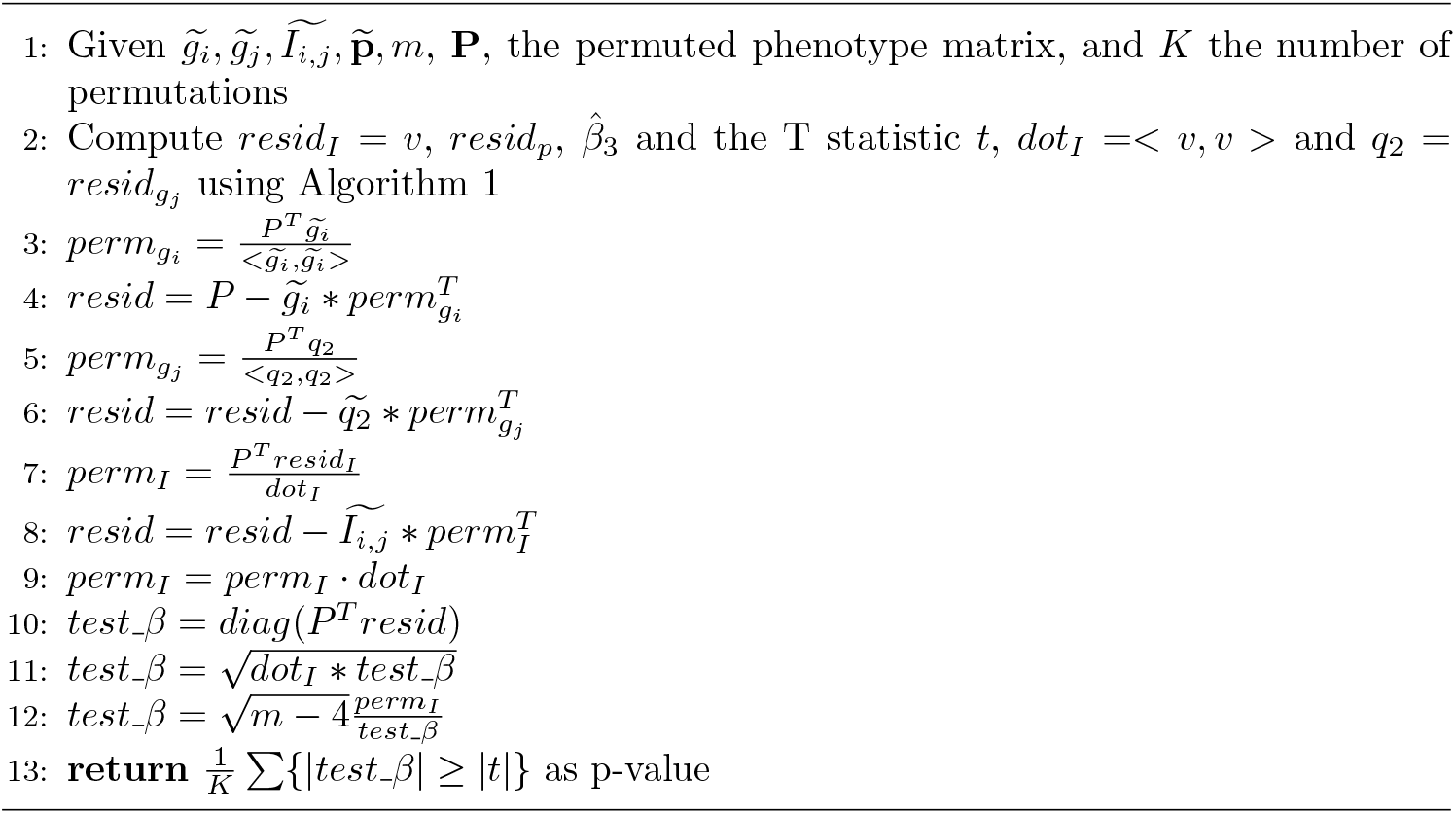

### Screening for higher order epistasis

We consider whether there is any mathematical justification from the models for epistasis to prune the search space of higher order interactions from lower order interactions or vice versa. For example, a common heuristic is to search the main effects for the pairwise epistasis and pairwise epistasis for 3-way epistasis. A variant of an approach that uses stages like this is used for example by Laurie et al [47].

Without loss of generality we are considering *k* + 1 loci, *g*_1_, … *g*_*k*+1_ and their interactions are all encoded using Cartesian (product) encoding. If such justification exists we may be able to show claims such that if there is a significant *k* + 1-way interaction among *k* + 1 loci, then all *k* interactions in the set of *k* + 1 loci are significant. The contrapositive of this claim is more practically useful: If any *k*-wise interaction is not significant from the set of loci, then *k* + 1-wise interaction is also not significant. The converse (i.e. if for a set of *k* + 1 loci, all *k*− wise epistasis exist, then the loci have *k* + 1 wise epistasis) is less plausible still. There is empirical evidence in the rat data set in this study (and many others) against both claims in considering only main effects and pairwise epistasis. Namely, there are main effects loci that are not in pairwise epistasis with each other; there are main effect loci in pairwise epistasis with a loci that is not a main effect and there exists pairwise epistasis between loci that are not main effects.

Nevertheless we examine why these claims do not hold for the models used in epistasis. To do so we consider two loci and their staged models for main effects and pairwise epistasis. Here we express the models simultaneously assuming the variables are centered. First we consider main effects:

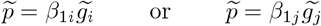

and pairwise epistasis

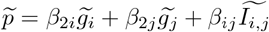

In terms of the coefficients in the models the first claim is that at least one loci not being a main effect implies no pairwise epistasis: if 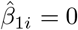 or 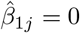, then 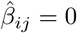. The concern here is really the inference needed regarding the coefficients of the pairwise model given knowledge of the coefficients in the main effects model. While having evidence 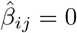 does not give us implications about 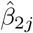 or *β*_2*i*_, knowing that either is 0 would give an implication regarding 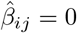. In general, without loss of generality, recent closed forms for the ordinary least squares coefficients [20] give extensions to the formula of Yule [45],

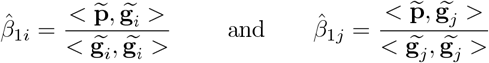

and for the pairwise epistasis model

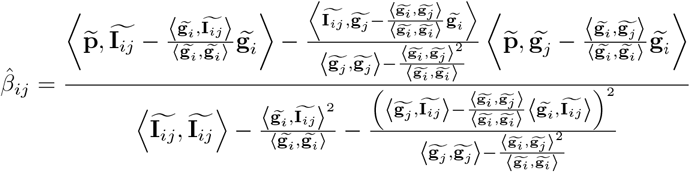

In the numerator for 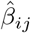, we note there are components that may be expressed in terms of the main effect coefficients, 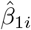 and 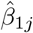. These will be the following two components:

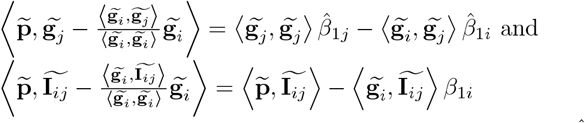

As a result, in terms of the main effect coefficients, 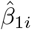 and 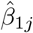, we may write the pairwise epistasis interaction term as

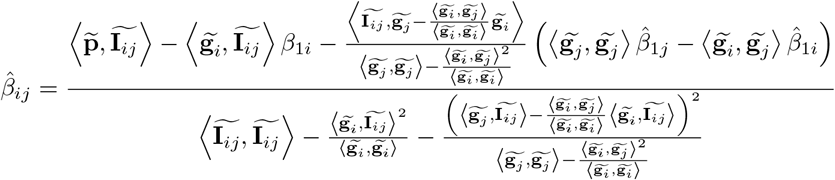

From this we observe that the numerator of 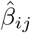 is determined upon the numerical value of the product 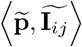 and zero values for 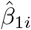 and 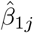 do not imply 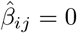. So insignificant interaction term, or value of 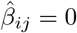 does not imply 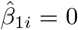 or 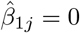 and on the other hand 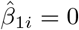 or 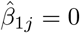 does not imply 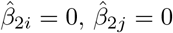 or 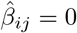. This may be considered a warning that we may not be justified only searching main effects for pairwise epistasis. Rather a biological reason for pruning loci, such as pruning of SNPs in LD, for epistasis may be more justified. For example, in the rat data set we applied the epistasis algorithms to loci screened from a previous GWAS study because of biological interest in those loci for the phenotype of BMI [21–23]. In a recent study in yeast, Ang et al. observed that epistasis most often occurred between loci with minor allele frequency between five and ten percent [48], so pruning loci for epistasis based on minor allele frequency may be another strategy for pruning the search space for epistasis detection for biological reasons.

### Application to Body Mass Index Data

We have applied our algorithm to a dataset of an outbred, related rat (*Rattus norvegicus*) population of males and females derived from eight inbred founders (Heterogenous Stock, [49]) in order to detect possible epistatic interactions associated with Body Mass Index (BMI). We also use the rat data to showcase the extension of our algorithm to higher order epistasis by investigating three-way interactions between putative QTLs identified via GWAS [21–23]. Additionally, we have applied our algorithm to a dataset of an inbred population of mice (*Mus musculus*) derived from 17 mouse strains [24, 25] to compare epistatic interactions, both Cartesian and XOR, in a closely related species to rat.

### Detecting Epistasis

The GWAS dataset of body mass index (BMI) residuals (corrected for sex and location) in 5,566 rats (*Rattus norvegicus*) containing approximately 129,000 autosomal SNPs was used for our rat analysis [21–23]. We selected the top 10,000 SNPs (SNPs with the 10,000 lowest *p*-values) from the original GWAS for further analysis as related work has illustrated positive correlations between main effects of a loci and instances of detected epistasis [7, 8]. For mice, we used the genotype/phenotype information from the Wellcome Trust Mouse Genomes Project [24, 25] found in the BGLR package [50] in R [51]. We extracted the genotype and BMI phenotype data available for 10,347 SNPs for 1,814 mice.

To account for the relatedness of the samples and population structures, the genetic relatedness matrices (GRM) were calculated for the rat data using the method of Yang et al. using the GCTA software tool [52] and the mouse data using the method of Sul et al [53]. We used the GRMs to calculate the variance component analysis for the phenotype of BMI for each data set using the methods of Joo et al. [54] and Kang et al. [55, 56] as implemented in the mmer function with method EMMA of the sommer package version 3.2 in R [57]. From the variance component analyses we obtained the inverses of the covariance matrices for both data sets as returned by the sommer package and then used the the sqrtm function from the expm package version 0.999-7 in R to obtain their square roots [58]. These inverse square root covariance matrices were then used to correct for population structure and essentially solve a generalized linear model by performing a weighted least squares by multiplying each variable in the model by the half inverse matrix before applying our algorithms as a pre-processing step. This “mixed model trick” is summarized by Suh et al [53].

Both Algorithms 1 and 2 were implemented in Python 3.9 for our experiments. To calculate the *p*-values for the two-sided T tests for the test statistics returned from our algorithms, the t.sf function from the scipy.stats package was used [59]. For two-way epistasis, Algorithm 1 was applied to all possible pairs of 10,000 SNPs in the rat data set and all possible pairs of 10,347 SNPs in the mouse data set. This was done as two separate experiments for each data set, once for the XOR and another for the Cartesian product. In addition for the XOR penetrance function for all 10,000 SNPs for the rat data set, *p*-values were also calculated using the permutation testing algorithm with 1000 permutations. (This was computationally intensive since we calculate nearly fifty billion tests and was done in parallel.) To account for multiple testing, FDR as implemented in the fdrcorrection function in the statsmodels package in Python [60] using a *p*-value threshold of 0.05 was applied. For three-way epistasis for the rat data set, we applied Algorithm 2 to the 18 putative main effect QTLS from the rat GWAS study [21–23] to see if any were involved in significant three-way epistatic interactions.

### Pruning Redundant Epistatic Events

For mice, we map all SNPs to their respective genomic location (assembly GRCm39) and retain those that map to the 19 autosomes in mice. This leaves us with 9,525 SNPs for further analyses. Although GWAS SNPs were LD-pruned, it is likely that many detected pairs are redundant in that one or both epistatic partners of a particular pair are in LD with others in close genomic proximity. We choose a conservative threshold of 10Mb upstream or downstream to prune redundant pairs in both species. All pruning steps are performed in R [51]. Under both Cartesian and XOR encodings to prune interchromosomal pairs, four conditions must be met: 1.) locus one in pair one and locus one in pair two are on the same chromosome, 2.) locus two in pair one and locus two in pair two are on the same chromosome, 3.) the absolute value of the difference in chromosomal position (in bp) between locus one in pair one and locus one in pair two is less than 10Mb, and 4.) the absolute value of the difference in chromosomal position between locus two in pair one and locus two in pair two is less than 10Mb. If all four conditions are true, the epistatic pair with the lower FDR-corrected *p*-value is retained while the other is omitted. We then check for mirror redundancies where two pairs technically meet the above criteria, but the chromosomal combination is reversed. The pair that is identified first is retained while the other is omitted. To prune intrachromosomal pairs, we check if two pairs exist on the same chromosome and then use the same bp criteria as outlined above for interchromosomal pairs. We also take an additional pruning step for intrachromosomal pairs where the absolute value of the bp difference between locus one and locus two is less than 10Mb. If this condition is met, the pair is omitted. This step removes close-acting cis-regulatory epistatic events as a byproduct of controlling for high LD.

There are nine total multi-locus genotypes (MLGs) possible under pair-wise epistasis of two biallelic SNPs (File S2). We limit our results to only those pairs where all nine MLGs occur. Additionally, we remove any significant epistatic pair, both in Cartesian and XOR, that do not have at least 10 or 5 observations of all nine MLGs in the datasets in rats and mice, respectively. Under Hardy-Weinberg assumptions, if the minor allele frequencies in two loci are 0.10, then we only expect to observe a genotype frequency for a double minor homozygote of 0.0001. We select the cutoff observations of 10 and 5 in rats and mice, respectively to ensure the combinatory genotype frequencies of the MLGs to be sufficiently above what would be expected if minor allele frequencies are 0.10 in both epistatic loci under Hardy-Weinberg assumptions.

### Identifying QTL-Associated and non-QTL-Associated Epistatic Events and Epistatic Hubs

After our initial pruning steps, we determine if a locus was associated with a putative QTL for BMI (“BMI with tail” in rats and “BMI” in mice) from the original GWAS studies [21–25] to observe if most epistatic events occur at or near loci with large main effects. To accomplish this, we use lists of the putative single locus QTL from the original GWAS studies and their respective genomic locations. We record if either locus 1 or locus 2 of a pair is within 10Mb upstream or downstream of a putative QTL to count the number of epistatic pairs with one putative GWAS QTL involved. We also note if both loci of a pair are associated with a GWAS QTL to record QTL to QTL epistatic interactions. If a locus is within 10Mb upstream or downstream of a GWAS QTL, its identification is replaced by the putative QTL’s identification. We then count how many epistatic events involve each GWAS QTL. If non-QTL-associated loci are within 10Mb upstream or downstream of each other on the same chromosome, the average genomic location is calculated and all non-QTL epistatic loci’s bp positions used to calculate the average are replaced by the average bp location. This procedure identifies non-QTL-associated epistatic loci. For our analyses, any locus with 10 or more epistatic interactions are defined as an epistatic hub (GWAS QTLs included).

Under XOR, we also identify loci that are within 10Mb upstream or downstream of a GWAS QTL and replace their identifications with the respective GWAS QTL. For non-QTL-associated XOR loci, we determine which loci are within 10Mb upstream or downstream of the non-QTL-associated loci identified under Cartesian encoding and replace their identification with the identification of the Cartesian locus. This allows us to determine how many epistatic loci are shared between both encodings. For loci that are unique to XOR encoding, we apply the same procedure used to determine non-QTL-associated loci under Cartesian encoding where we calculate the average genomic location among loci if on the same chromosome and within 10Mb upstream or downstream of each other. Thus, we identify loci specific to each encoding as well as loci shared between encodings.

### Gene Set Enrichment and Kegg Pathway Analysis

To perform functional annotation for loci involved in epistatic events, we query protein-coding gene models, non-protein coding gene models, and pseudogene models from the Rat Genome Database (https://rgd.mcw.edu/) for rats and mice. We retrieve any model that is 1Mb upstream or downstream from each epistatic locus. We then convert the species-specific gene model symbols from the respective database to Entrez IDs. Any returned queries that do not have associated Entrez IDs are omitted. We use this list to identify the enrichment of GO (gene ontology) terms (cellular component, molecular function, and biological process) using the BioConductor ([61]) package clusterProfiler [62, 63] in R with FDR correction (*p*-value cutoff= 0.01; *q* -value cutoff= 0.05). We also perform KEGG pathway enrichment analysis using clusterpPofiler (*p*-value cutoff= 0.05; *q* -value cutoff= 0.02). Dotplot figures of enriched ontological terms are made using the DOSE [64] package in R.

### Identifying Three-Way Epistatic Interactions between Putative QTLs

To test the extension of our algorithm for identifying higher order epistasis, we explore three-way epistatic interactions among the 18 putative QTL identified in the original rat GWAS [21–23] under both Cartesian and XOR interaction encodings. Triplets that have an associated experimental *p*-value *<* 0.05 are retained and we record occurrences of three-way epistasis for each GWAS QTL. In addition, the triplets between each encoding (Cartesian or XOR) are compared for overlap. Since all significant triplets in both experiments are unique in terms of genomic location and between putative QTL, no pruning or epistatic locus identifications are performed for this analysis.

## Results

### Most Epistatic Interactions and Hubs Occur at non-QTL-Associated Loci

For the Cartesian experiment in rats, our method detects 4,158 (86.8%) interchromosomal and 634 (13.2%) intrachromosomal significant pairs (4,792 total; File S1). After MLG pruning, this reduces to 3,109 (90.4%) interchromosomal and 329 (9.6%) intrachromosomal significant pairs (3,438 total; File S1). After redundancy pruning, this is further reduced to 175 interchromosomal pairs (96.2%) and seven intrachromosomal pairs (3.8%) (182 total) (Fig 1A; File S1). There are 182 Cartesian pairs after all pruning. Of these, there are 66 pairs (36.3%) containing one QTL-associated locus and 9 (4.9%) QTL-QTL interactions (Fig 1C; File S1). However, most pairs are between two non-QTL-associated loci (107 pairs; 58.8%). There are a total of 91 unique Cartesian epistatic loci. Of these, 75 are non-QTL-associated (82.4%) (Fig 1E; File S1). Of the 18 putative GWAS QTL, chr7:8599340_A, chr18.32316331_A, and chr5:72916242_T are not involved in epistasis (File S1). It is important to note that chr18.32316331 A is physically close to another putative QTL, chr18.27348077_G (4,968,254 bp apart), and is not represented as loci we detect are physically closer to chr18.27348077_G.

**Fig 1.**
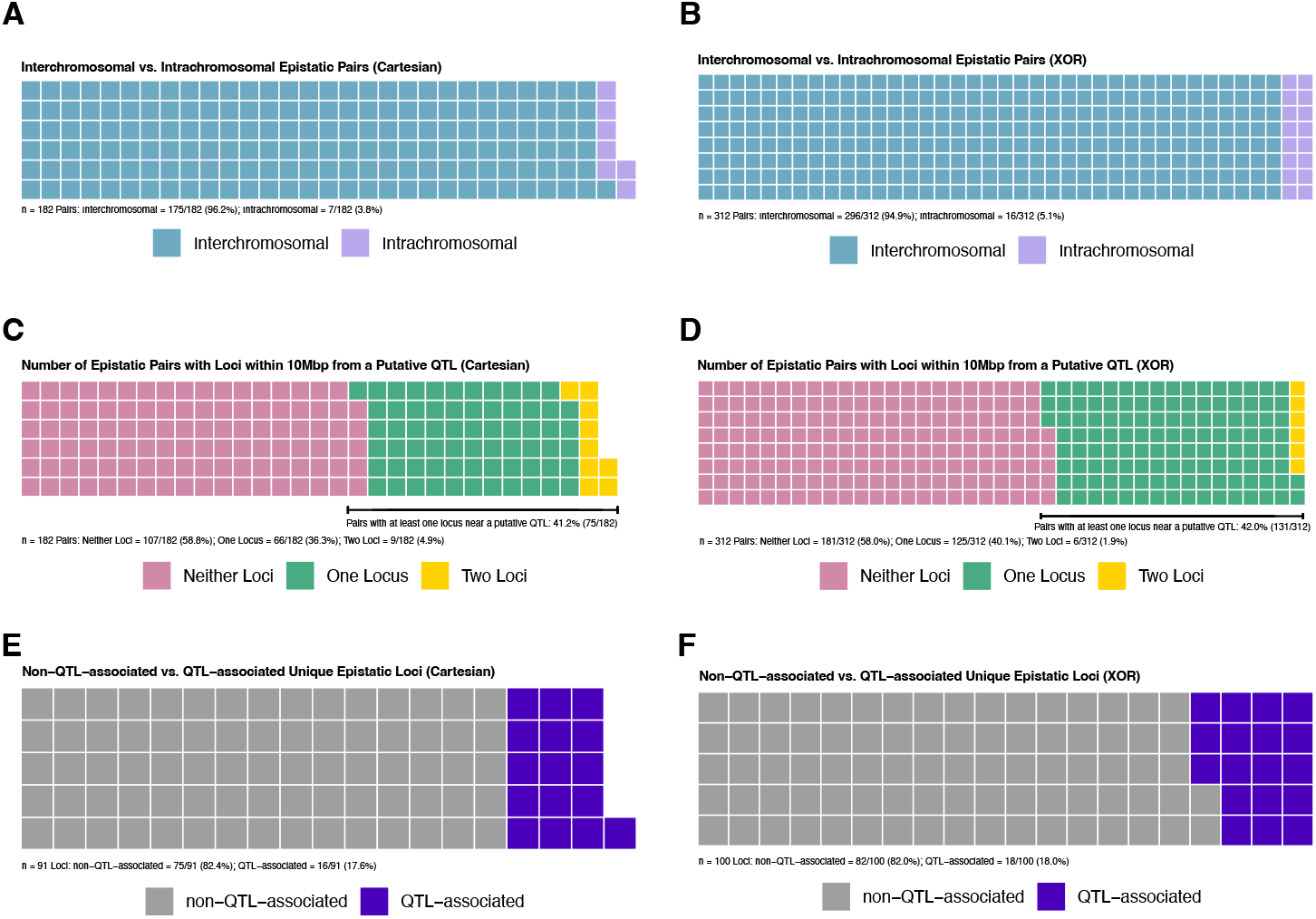
Proportions of epistasis detected in rats. A and B: Interchromosomal (blue squares) vs intrachromosomal (light purple squares) pairs under Cartesian (A) and XOR (B) models. C and D: Number of epistatic pairs involving no (pink squares), one (green squares), or two (yellow squares) putative GWAS QTL under Cartesian (C) and XOR (D) models. E and F: Non-QTL-associated (gray squares) and QTL-associated (dark purple squares) epistatic loci under Cartesian (E) and XOR (F) models.

In mice, 81 epistatic pairs are detected under Cartesian encoding. However, 65 of these involve at least one locus on the X chromosome and are omitted as we are only investigating autosomal loci, leaving 16 pairs. None of these pairs are intrachromosomal (File S3). After MLG pruning eight pairs remain and after redundancy pruning only one pair remains between chr3.44666611, and chr12.7079769 (File S3). In rats, XOR encoding yields more significant pairs compared to Cartesian (31,182 vs. 4,792) (File S1). Of the 31,182 significant pairs, 27,774 are interchromosomal (89.1%) and 3,408 are intrachromosomal (10.9%; File S1). After MLG pruning, this is reduced to 11,016 (86.4%) interchromosomal pairs and 1733 (13.6%) intrachromosomal pairs (12,749 pairs total; File S1). After redundancy pruning, this is further reduced to 296 (94.9%) interchromosomal pairs and 16 (5.1%) intrachromosomal pairs (312 pairs total) (Fig 1B File S1). Most pairs (181 (58.0%)) are between two non-QTL-associated loci while there are 125 (40.1%) pairs involving one QTL-associated locus and six (1.9%) QTL-QTL interactions (Fig 1D; File S1). Epistatic pairs involve 100 loci where 82 (82.0%) are non-QTL-associated (Fig 1F; File S1). All 18 putative GWAS QTL are represented under XOR encoding. In mice, under XOR, there are 468,596 significant pairs. After we map these loci to genomic locations and remove any pair involving an non-autosomal locus, 25,055 pairs remain with 23,688 (94.5%) interchromosomal and 1,367 (5.5%) intrachromosomal (File S3). After MLG pruning, this is reduced to 13,328 (95.0%) interchromosomal pairs and 707 (5.0%) intrachromosomal pairs (14,035 pairs total; File S3). Finally, after redundancy pruning, this is reduced to 341 (92.4%) interchromosomal pairs and 28 (7.6%) intrachromosomal pairs (369 pairs total) (Fig 2A; File S3). As in rats, and due to there only being 3 putative GWAS QTL for BMI in mice we were able to map to a genomic location, most pairs (351 (95.1%)) are between two non-QTL-associated loci while there are 18 (4.9%) pairs containing one QTL-associated locus (Fig 2B; File S1). There are no QTL-QTL interactions in mice. In mice, XOR yields 115 unique epistatic loci where 112 (97.4%) are non-QTL-associated.

**Fig 2.**
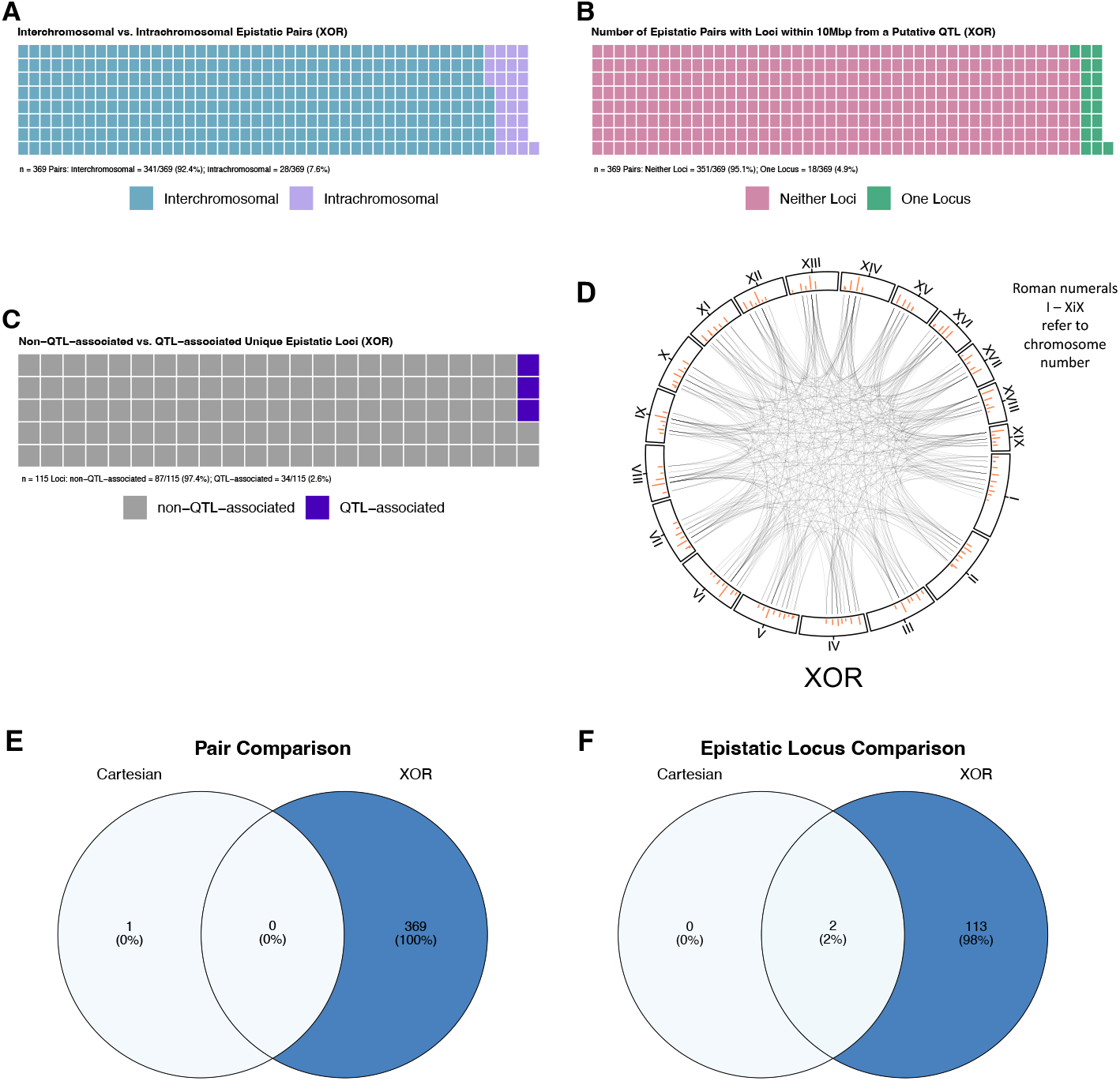
Representations of epistasis in mice. A: Interchromosomal (blue squares) vs. intrachromosomal (light purple squares) pairs under XOR model. B: Number of pairs involving no (pink squares), one (green squares), or two (yellow squares) putative GWAS QTL under XOR model. C: Non-QTL-associated (gray squares) and QTL-associated (dark purple squares) epistatic loci under XOR model. D: Interaction plot of XOR epistasis across mouse autosomes. Autosome numbers are depicted as roman numerals and increase clockwise. Orange bars represent counts of epistatic instances per locus. E: Venn diagram of epistatic pairs under Cartesian (left circle) model, the intersect between models (center), and under XOR (right circle) model. F: Venn diagram of epistatic loci under Cartesian (left circle) model, the intersect between models (center), and under XOR (right circle) model. E and F: Color gradient illustrates low (white) to high (blue) occurrences.

It is important to note that a large amount of significant epistatic pairs involve non-autosomal or unmappable loci in mice. In future analyses, we aim to explore interactions that involve non-autosomal loci within the mouse cohort and in other species.

The rank order and number of interactions per GWAS QTL differ between experiments and across species. In rats, under Cartesian encoding, the largest QTL-associated hubs are chr1.281788173_G and chr5.107167969_G with 12 interactions each (Fig 3A; File S1). chr1.281788173_G is also the locus with the largest main effect signal in the rat GWAS (File S1). Under XOR encoding, there are four QTL-associated hubs identified (Fig 3B; File S1). As in Cartesian, chr1.281788173_G is the largest QTL-associated hub under XOR with 34 interactions. This is followed by chr18:27348077_G with 25 interactions. XOR QTL-associated hubs are also identified at chr5:107167969_G, and chr8:103608382_G. Under Cartesian encoding, 10 of the putative GWAS QTLs are in QTL-QTL interactions (Fig 3C; File S1). Two putative GWAS QTLs are in three QTL-QTL interactions while four are in two QTL-QTL interactions The remaining four are in one. The Cartesian QTL hubs, chr5.107167969_G and chr1.281788173_G are involved in QTL-QTL interactions, but there doesn’t seem to be a clear relationship between number of epistatic interactions and number of QTL-QTL interactions. Under XOR encoding, seven putative GWAS QTL are in QTL-QTL interactions (Fig 3D; File S1). However, the rank orders and QTL representations are distinct between encodings. For example, under XOR, chr1.281788173_G has the largest occurrences of QTL-QTL interactions with five. Under XOR, there also is not a clear relationship between the number of QTL epistatic events a hub is involved in and the propensity of QTL-QTL interactions.

**Fig 3.**
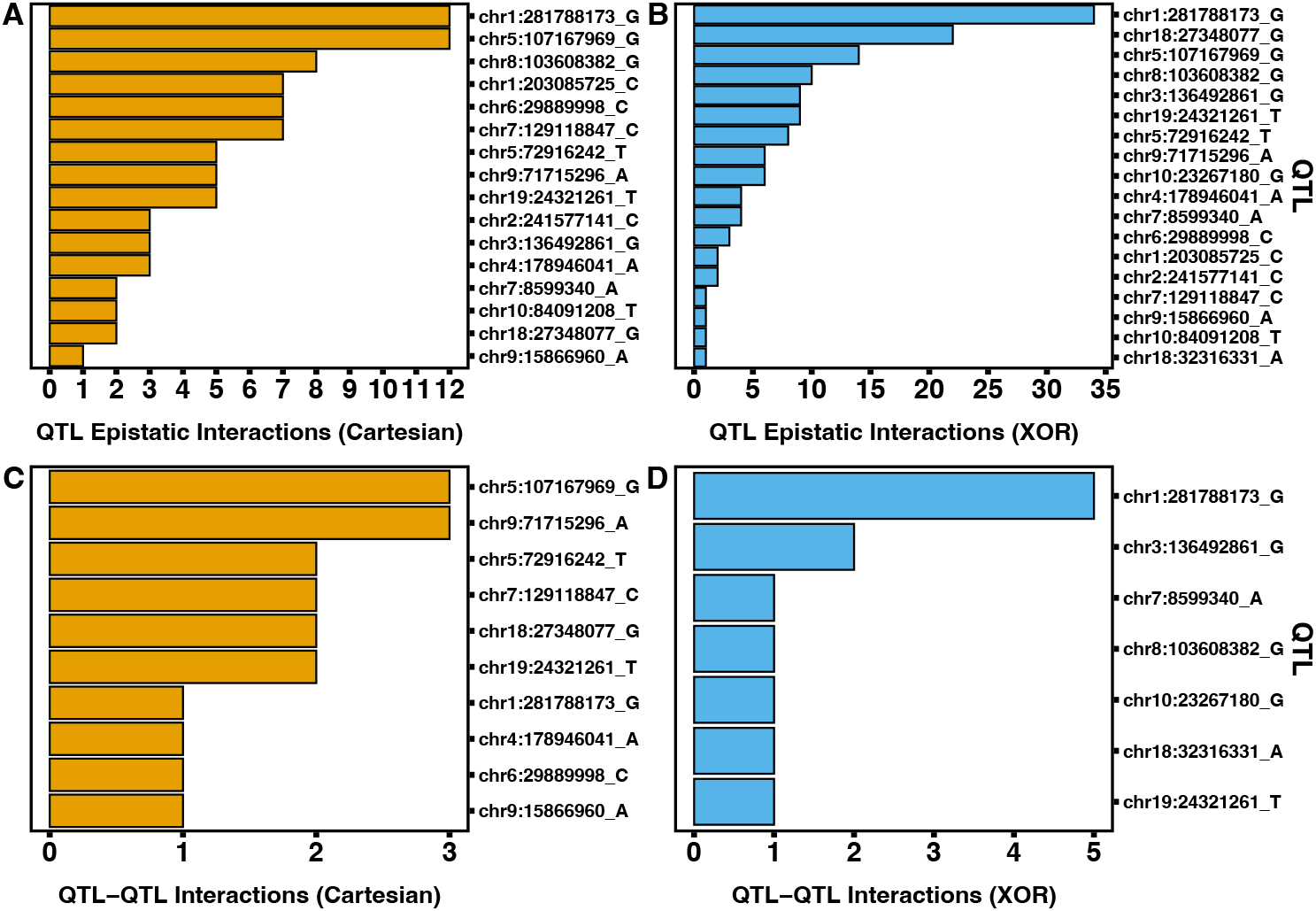
Bar graphs of occurrences of QTL-associated epistasis in rats. A and B: Bar graphs of instances of epistatic events for GWAS putative QTL under Cartesian (A, orange bars) and XOR (B, blue bars) models. C and D: Bar graphs of instances of QTL to QTL epistatic events for GWAS putative QTL under Cartesian (C, orange bars) and XOR (D, blue bars) models.

In mice under Cartesian encoding, QTL and QTL-QTL interactions do not occur as only one significant pair, involving two non-QTL-associated loci, is detected. Under XOR encoding, all three putative QTL are involved in a least one epistatic interaction however none are hubs (Fig 4A; File S3). The largest QTL epistatic loci is chr2.68831331_G with eight interactions. No QTL-QTL interactions occur in mice under XOR encoding.

**Fig 4.**
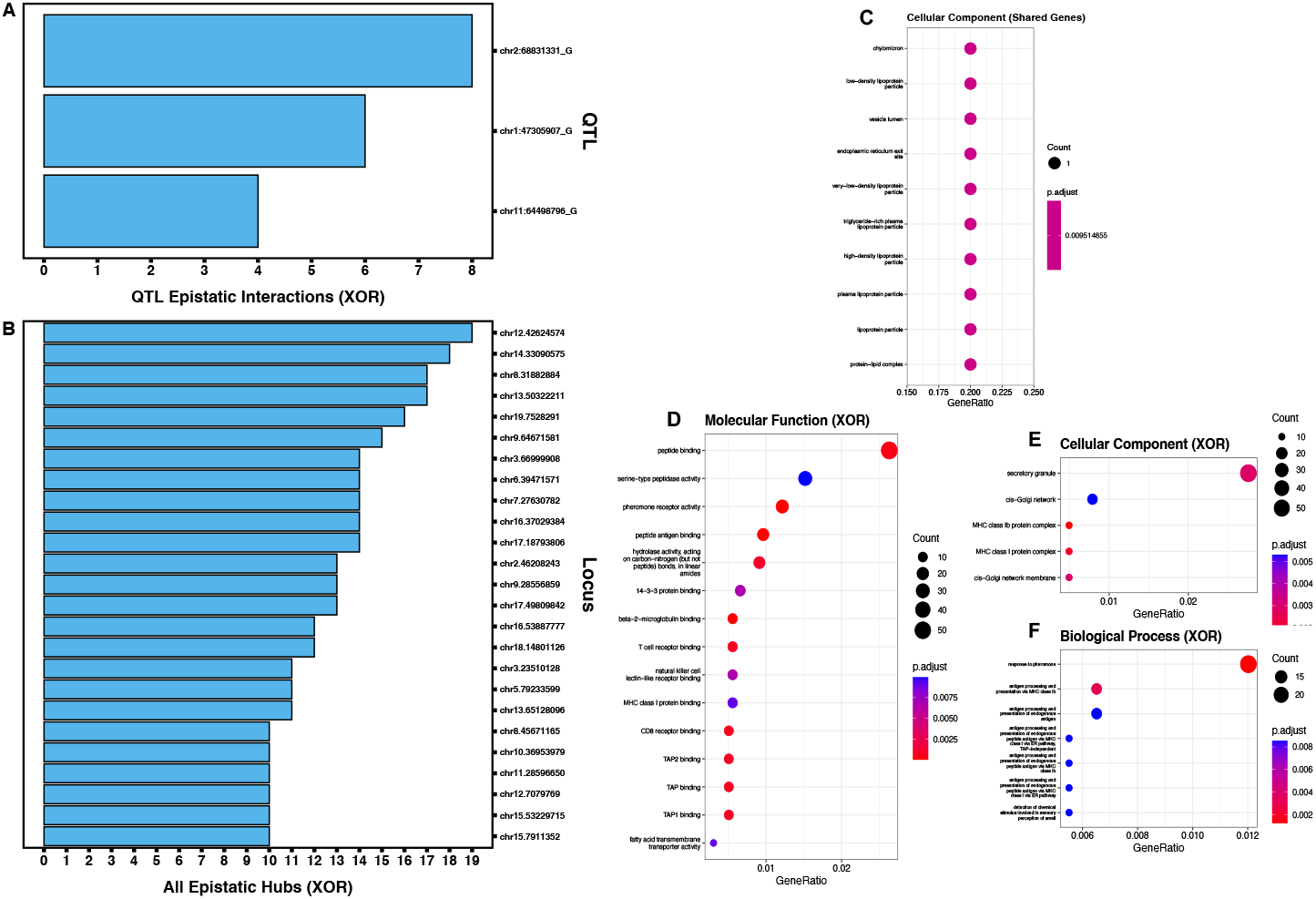
Bar graphs of epistatic interactions and dot plots of GO term gene set enrichment in mice under XOR encoding. A: Bar graph of instances of QTL-associated epistatic events for GWAS putative QTL under XOR model. B: Bar graph of instances of epistasis for all epistatic hubs (QTL-associated and non-QTL-associated) under XOR model. C: dot plots for enriched cellular component GO terms. D through F: dot plots for enriched GO terms for biological processes (D), cellular components (E), and molecular functions (F) for XOR-specific epistatic loci. C through E: size of dot corresponds to the number of genes associated with that enrichment. Color gradient illustrates level of significance with higher *p*-values in blue and lower *p*-values in red.

In rats, there are four Cartesian non-QTL-associated epistatic hubs under Cartesian and 18 under XOR (Fig 5A,C; File S1). The largest non-QTL-associated epistatic hubs are chr16.54146824 under Cartesian and chr9.60993465_G under XOR. As we observe with GWAS QTL epistasis, the rank order and hub sizes of non-QTL-associated hubs differ between experiments (Fig 5A,C; File S1). When including putative GWAS QTLs as hubs, there are a total of six epistatic hubs identified under Cartesian encoding (Fig 5B; File S1) and 22 under XOR encoding (Fig 5D; File S1). GWAS QTLs account for 33% (2/6) and 18.2% (4/22) of Cartesian and XOR hubs, respectively. Despite the rank order of hubs being largely distinct between experiments, the largest hub under both Cartesian and XOR encodings is the putative GWAS QTL with the largest signal in the rat GWAS [21–23], chr1.281788173_G. In mice, under XOR encoding, all 25 hubs are non-QTL-associated. The largest is chr12.42624574 with 19 interactions (Fig 4B, File S3).

**Fig 5.**
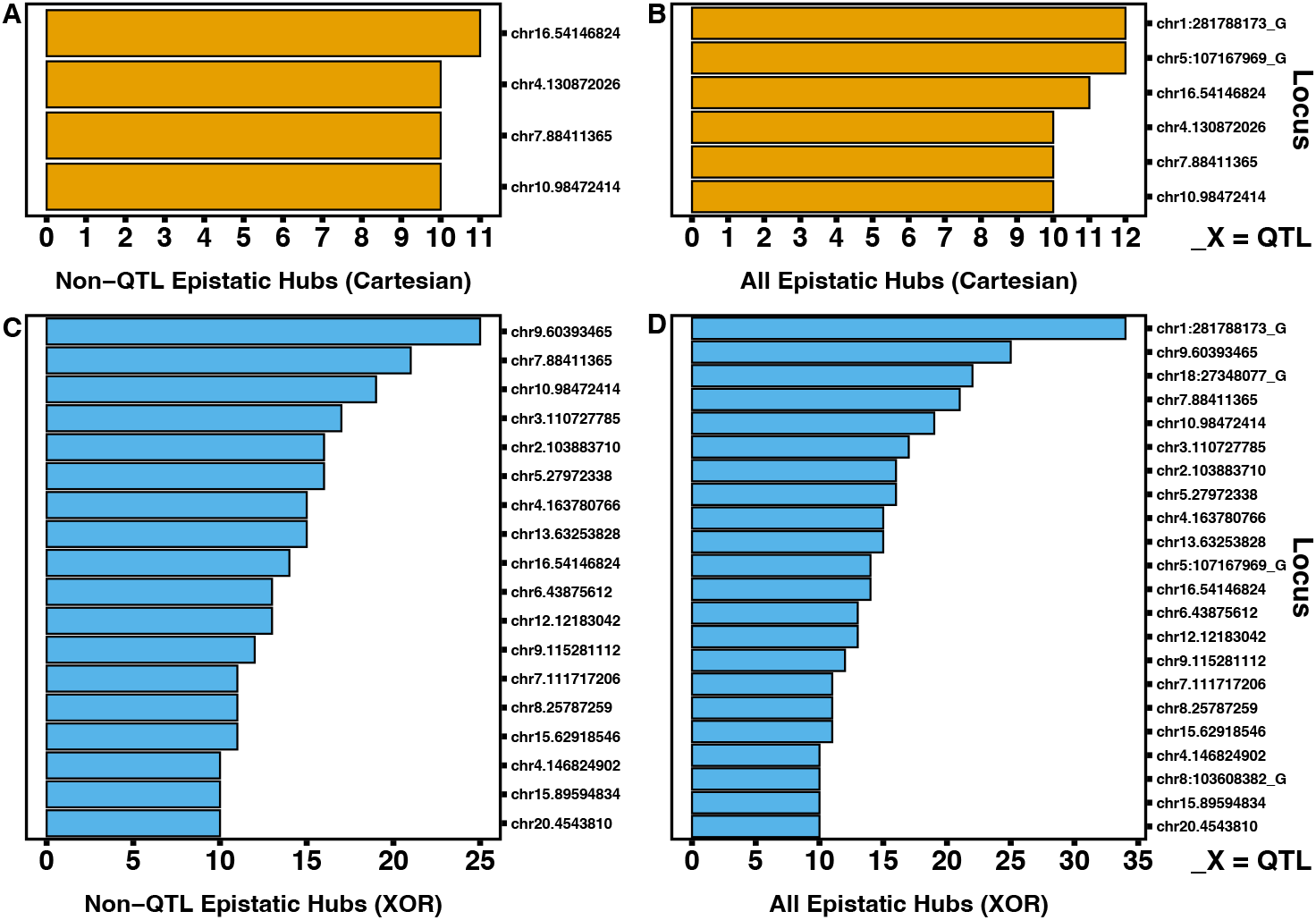
Bar graphs of epistatic events in non-QTL-associated hubs and all hubs in rats. A and B: Bar graphs of instances of epistatic events for non-QTL-associated hubs (A) and all hubs (B) under Cartesian model (orange bars). C and D: Bar graphs of instances of epistatic events for non-QTL-associated hubs (C) and all hubs (D) under XOR model (blue) bars). B and D: Loci on y axes ending with “_“ and allele designation are GWAS putative QTL.

In rats, there is a significant correlation between the significance of SNPs in the GWAS considering its *p*-value (i.e. -log10*p*-value) and the number of epistatic interactions under XOR encoding when considering all epistatic SNPs (*p* = 0.0112, r = 0.253; File S2) and only QTL-associated SNPs (*p* = 0.00249, r = 0.667; File S2). This relationship does not occur under Cartesian encoding. We did not perform this analysis in mice as we did not have access to the GWAS statistics for all SNPs.

Permutation test results in rats under XOR encoding yielded 29 XOR specific loci compared to 25 when just using FDR correction alone. Of the 29 XOR specific loci after 1000 permutations, most (23) overlap (are within 1 Mb of a similar locus with FDR correction alone) while six are only found after 1000 permutations (File S1). In this instance permutation testing leads to the detection of additional epistatic loci. We only apply the permutations algorithm to epistatic pairs from rat data under XOR encoding to both test the algorithm’s capability and to verify detection of XOR statistical epistasis in a living system.

### Cartesian and XOR Share Common Epistatic Loci while Epistatic Landscapes are Distinct

In rats, the interaction landscapes of Cartesian and XOR two-way epistasis are mostly distinct (Fig 6A,B; File S1). Out of the 182 Cartesian and 312 XOR pairs, only 16 pairs (3%) occur under both encodings (Fig 6C; File S1). However, of the 91 and 100 epistatic loci that occur under Cartesian and XOR encodings, respectively, 75 (65%) are shared Fig 6D; File S1). Although most epistatic loci are shared between Cartesian and XOR, distinct significant two-way epistatic interactions occur under each encoding.

**Fig 6.**
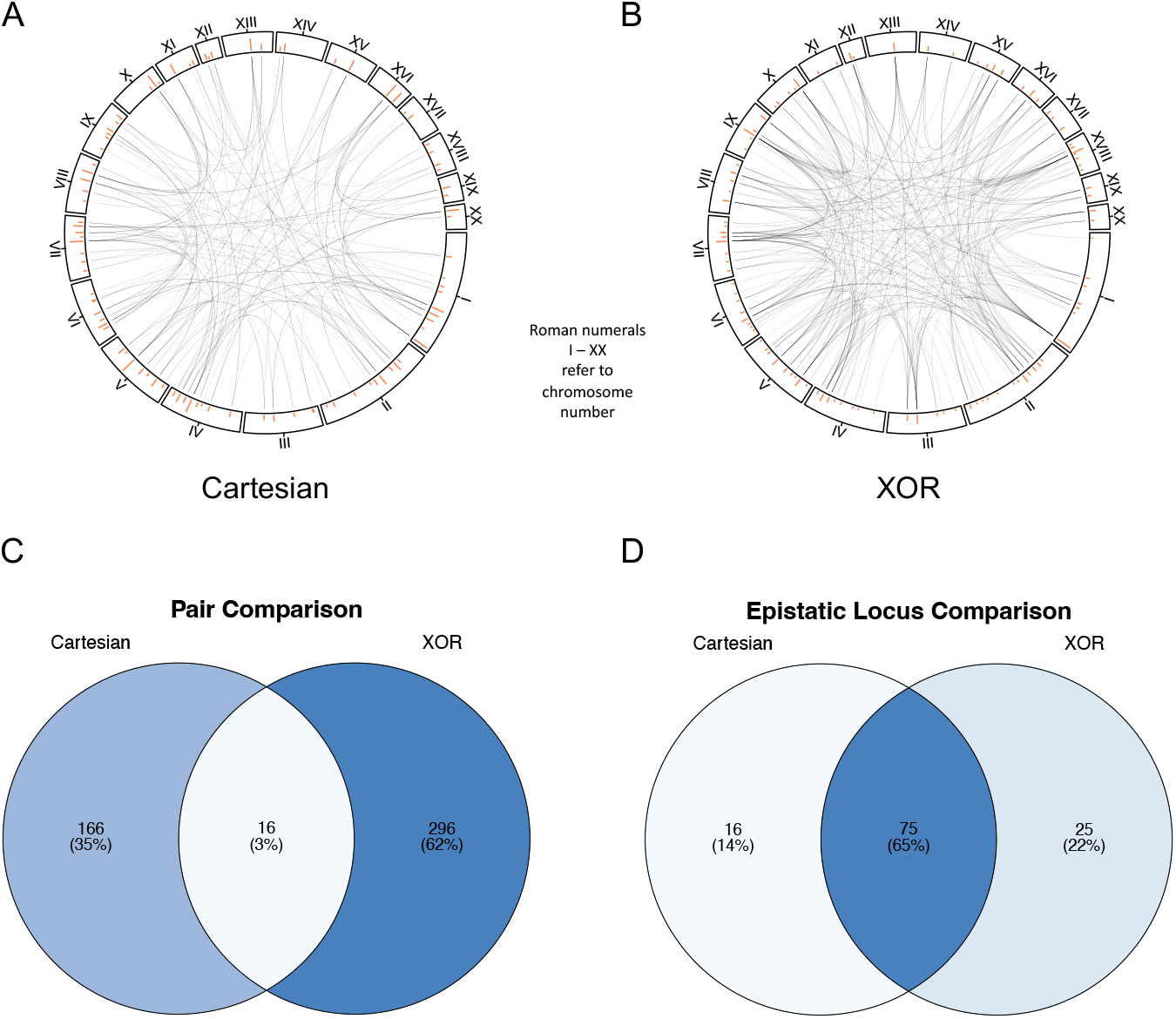
Interaction plots and epistatic model comparisons in rats. A and B: Interaction plots of Cartesian (A) XOR (B) epistasis across rat autosomes. Autosome numbers are depicted as roman numerals and increase clockwise. Orange bars represent counts of epistatic instances per locus. C: Venn diagram of epistatic pairs under Cartesian (left circle) model, the intersect between models (center) and under XOR (right circle) model. D: Venn diagram of epistatic loci under Cartesian (left circle) model, the intersect between models (center) and under XOR (right circle) model. C and D: Color gradient illustrates low (white) to high (blue) occurrences.

In mice, only one epistatic pair is detected and is distinct to Cartesian encoding (i.e., not detected using an XOR encoding) (Fig 2E; File S3). Only two epistatic loci are detected under Cartesian encoding. These two loci are also detected under XOR encoding (Fig 2F; File S3).

### Enriched Terms and Pathways Detected for XOR-Specific Epistatic Loci

In rats, the 16 Cartesian-unique and 25 XOR-unique epistatic loci are analyzed using gene set enrichment. Additionally, the 75 loci that are shared between the encodings and the 18 putative QTLs are also analyzed. In Cartesian encoding, the 16 Cartesian-unique loci led to no significant enrichment of cellular component, biological process, or molecular function. However, one KEGG pathway was enriched - Ribosome (File S1). Enrichment of the 75 shared loci reveal biological processes, cellular components, and molecular functions associated largely with immunity (Fig 7A-C; File S2). Under XOR encoding, five enriched biological processes are identified, all of which are metabolic in nature (Fig 7D; File S1; File S2). A total of 10 molecular functions are enriched with the vast majority being oxidoreductase activities and carboxylic acid binding (Fig 7E; File S1; File S2). No cellular components are significantly enriched. A total of 17 KEGG pathways are enriched. Notable metabolism-associated KEGG pathways are nitrogen metabolism, gastric acid secretion, and glucagon signaling pathway (File S1). In mice, the two shared loci between encodings reveal significant enrichments of metabolic terms (Fig 4C; File S3). The enrichments from the 113 XOR-specific loci are mostly involved in immunity (Fig 4D,E; File S3). There are a total of eight biological functions, five cellular components, 15 molecular functions, and two KEGG pathways significantly enriched from XOR-specific loci. However, “detection of chemical stimulus involved in sensory perception of smell”, “fatty acid transmembrane transporter activity”, and “hydrolase activity, acting on carbon-nitrogen (but not peptide) bonds, in linear amides” are enriched metabolically-relevant processes and functions. Additionally, both KEGG pathways are metabolic in nature (File S3).

**Fig 7.**
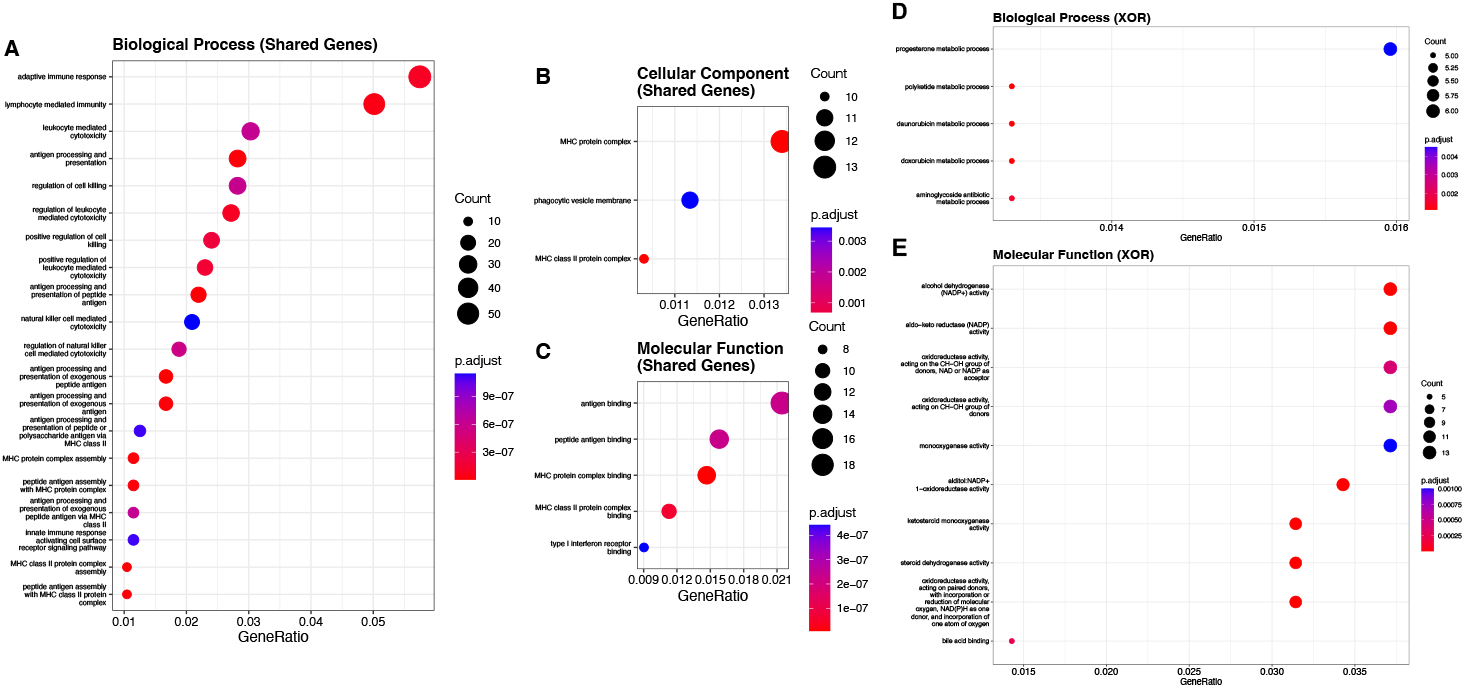
Dotplots of gene set enrichment in rats. A through C: dot plots for enriched GO terms for biological processes (A), cellular components (B), and molecular functions (C) for shared genes between Cartesian and XOR models. D and E: dot plots for enriched GO terms for biological processes (D) and molecular functions (E) for XOR-specific epistatic loci. A through E: size of dot corresponds to the number of genes associated with that enrichment. Color gradient illustrates level of significance with higher *p*-values in blue and lower *p*-values in red.

In rats, GO term enrichments for the 75 shared epistatic loci between encodings yielded 75 significantly enriched biological processes, three cellular components, five molecular functions, and 46 KEGG pathways. The vast majority of these terms and pathways are involved in immunity (File S1). In mice, the two shared epistatic loci between Cartesian and XOR encodings significantly enriched 10 cellular components and four KEGG pathways, all of which are involved in metabolism (File S3). This is likely due to the gene *ApoB* being located in close proximity to the chr12.7079769 epistatic locus. Interestingly, SNPs near this gene were not implicated in the original GWAS study ([24, 25]; File S3). In rats the 18 putative GWAS QTL yield 14 significantly enriched biological processes and one molecular function (File S1). Outside of “cellular response to alcohol”, all other enriched biological processes are related to immunity. The lone enriched molecular function is “type I interferon receptor binding” The 18 putative GWAS QTL yield KEGG pathway enrichment mostly associated with immunity. However, metabolic-related pathways like “alcoholic liver disease” and “lipid and atherosclerosis” are also enriched (File S1). In mice, the three putative GWAS QTL are enriched for one cellular component (“inner dynein arm”) and two metabolic KEGG pathways: “Glycoaminoglycan biosynthesis” and “Cholesterol metabolism” (File S3).

### Three-Way Epistatic Landscapes for QTL are Distinct Between Encodings

XOR encoding of significant three-way epistatic interactions between putative GWAS QTL results in more and larger epistatic hubs compared to Cartesian encoding (seven GWAS QTL hubs in Cartesian vs. all 18 GWAS QTLs in XOR) (Fig 8A,B; File S1). Additionally, as we also observe in the two-way experiments, the epistatic landscapes and rank order of epistatic loci are distinct between the two encodings (Fig 8A-E; File S1). Furthermore, out of the 51 significant epistatic triplets in Cartesian and 90 in XOR, only 4 triplets are shared between them despite sharing all epistatic loci (the 18 putative GWAS QTLs) (Fig 8E; File S1). A similar result occurred in the two-way experiment (Fig 6D; File S1). There is no significant correlation between the absolute value of the GWAS betas or the GWAS -log*P* and the number of three-way interactions for the 18 putative GWAS QTL under either encoding (File S2).

**Fig 8.**
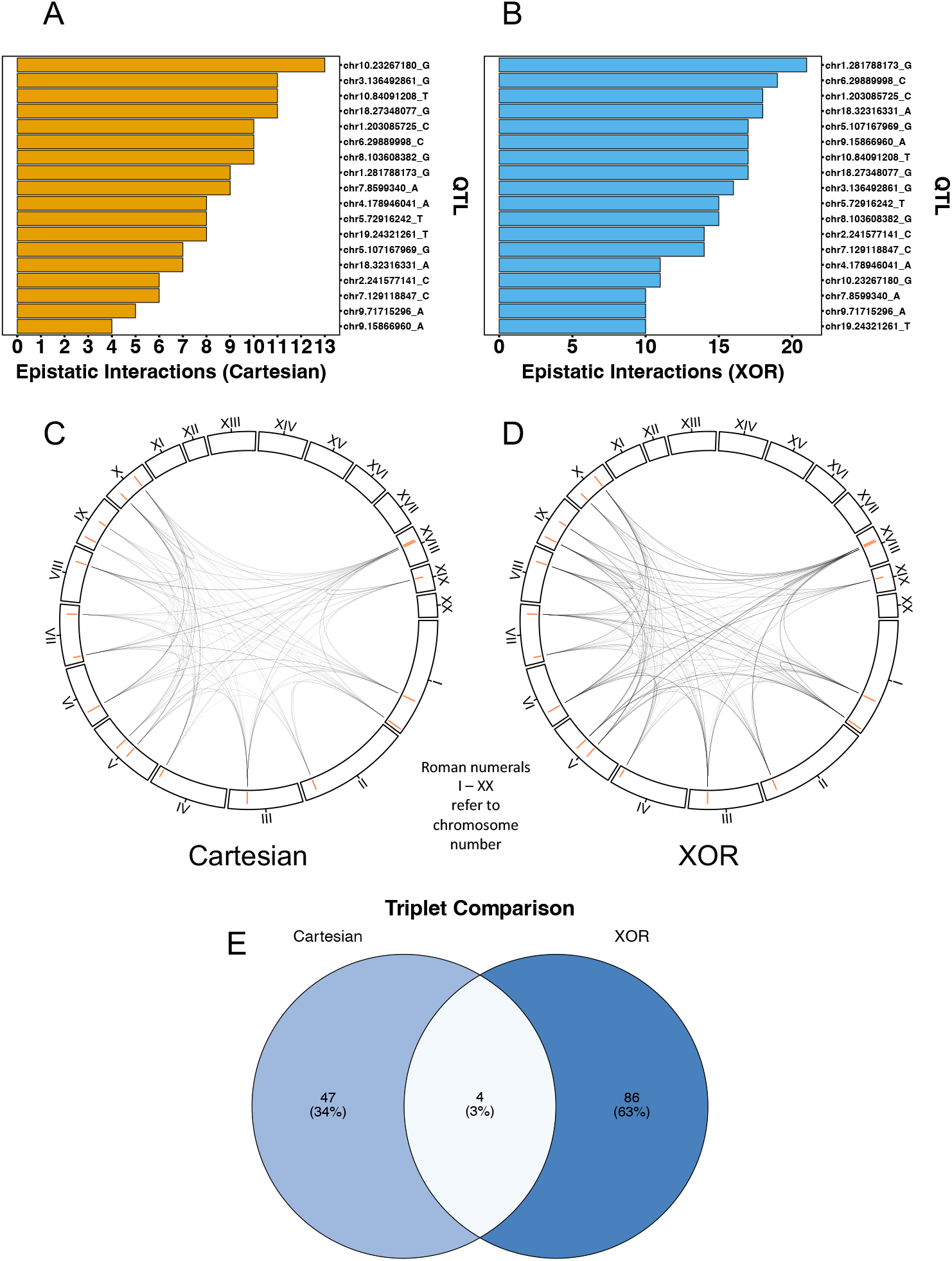
Representations of three-way epistasis in rats. A and B: Bar graphs of occurrences of three-way epistatic events in GWAS QTLs under Cartesian (A, orange bars) and XOR (B, blue bars) models. C and D: Interaction plots of Cartesian (C) XOR (D) three-way epistasis of GWAS QTLs across rat autosomes. Autosome numbers are depicted as roman numerals and increase clockwise. Orange bars represent counts of epistatic instances per locus. E: Venn diagram of epistatic triplets for GWAS QTLs under Cartesian (left circle) model, the intersect between models (center) and under XOR (right circle) model. E: Color gradient illustrates low (white) to high (blue) occurrences.

## Discussion

### Epistasis Occurs Widely at Non-QTL-Associated Locations

Our findings point to the potential ubiquity of epistasis in living systems as we have detected numerous epistatic pairs and loci under two distinct interaction encodings with many of them unique to either Cartesian or XOR. Furthermore, most of interactions detected occur at loci not associated with a GWAS QTL (more than 10Mb upstream or downstream) in both species. However, in rats, the largest epistatic hub under both Cartesian and XOR encoding is chr1.281788173_G, which was also the putative QTL with the largest signal found in the rat GWAS study. In mice, under XOR encoding, all epistatic hubs were non-QTL-associated. One explanation for this is that there are only three putative QTL for BMI from the mouse GWAS study that were mapped to a genomic location. Two studies investigating genome-wide non-additive effects in yeast found results to the contrary where a strong positive correlation between the main effect size of a locus and the number of interactions it was involved in is observed [7, 8].

Although we observed strong correlations in the two-way experiment between the GWAS p-value of a locus and number of interactions under an XOR encoding in rats, it is important to consider that most hubs, both under Cartesian and XOR encodings and in both species, are not located near a putative QTL. There are several possible explanations as to why most instances of detected epistasis occur at non-QTL-associated locations. The first is that there are more possible non-QTL-associated genomic locations that could serve as epistatic loci or hubs. The rat GWAS study identified 18 putative QTL. Therefore, only 360Mb of the *R. norvegicus* genome would be considered QTL-associated under our pruning and categorization strategy. The R. norvegicus genome assembly used in this study is approximately 2.6 Gb in size [65], meaning that there are more possibly non-QTL-associated regions we could have detected in this study compared to the 18 QTL-associated regions. The mouse GWAS identified four QTL for BMI, three of which mapped to a genomic location, representing only 30Mb of the mouse genome. This same reasoning as to why there are more non-QTL-associated epistatic loci can be applied to why we observe so many more interchromosomal epistatic pairs compared to intrachromosomal pairs. Simply, there are more possible pairwise combinations that can occur across chromosomes than can occur within the same chromosome. The second explanation is that main effects derived from significant GWAS summary statistics are not adequate predictors of epistasis. 82.4% and 82.0% of unique epistatic loci are non-QTL-associated under Cartesian and XOR encodings, respectively. Furthermore, the dataset of SNPs in mice were not selected based upon GWAS summary statistics. Yet, there were still significant levels of statistical epistasis detected. In this regard the mouse dataset is a more pertinent example of why searching for epistasis solely based upon univariate effects is not optimal. Even though, to test our methodology, we pruned the original rat dataset using GWAS p-values, we suggest alternative pruning strategies, perhaps based on expert knowledge or biological relevance, to test for epistasis.

Another alternative explanation is our pruning strategy for epistatic pairs itself. Our initial step is to remove redundant pairs within 10Mb upstream or downstream from each other and only the most significant pair (lowest p-value) is retained. In rats, there may be more redundant pairs involving loci in LD with putative QTL compared to non-QTL-associated pairs because our dataset is derived from the 10,000 most significant loci in terms of GWAS p-values. Perhaps with a different pruning strategy, QTL-associated loci would have possibly been highlighted more. However, our pruning strategy may highlight other loci, with non-significant main effects, as centers for epistasis. Taken together, our results highlight the importance of loci with non-significant main effects. It is plausible that combinatory mutations in multiple loci interacting in networks or pathways, may be required to explain much of the variation observed in phenotypes as canalized systems are likely resistant to alterations to one or few loci [16–19]. It is possible that we have detected loci underlying BMI in *R. norvegicus* and *M. musculus* that would otherwise go undetected if only main effects were considered. Additionally, the sheer number of loci involved in significant two-way interactions compared to the number of loci with significant main effects detected in the original GWAS point to the potential importance of epistasis in complex biological traits like BMI in two closely-related species.

### XOR Model Uncovers Numerous Epistatic Events not Detected under Cartesian Encoding

Although the number of epistatic loci detected in both Cartesian and XOR encodings is comparable and generally overlaps in rats, there are more significant pairs detected under XOR encoding compared to Cartesian. These pairings are mostly distinct with only 16 pairs shared between the encodings. In mice, there is only one significant Cartesian pair detected. This highlights that using a Cartesian encoding, at least in mice, may not uncover any substantial level of epistasis. If the XOR encoding for interactions had not been applied, nearly all of the detected epistasis in mice would have been missed. Furthermore, in rats, XOR hubs are larger and more numerous than Cartesian hubs. We also observe similar results in our 3-way experiment in rats with XOR hubs being larger and triplets being mostly unique between encodings. Taken together, these results suggest that interaction encoding is an important consideration when investigating epistatic events in biological systems.

The high amount of overlap between epistatic loci is partly attributable to re-using Cartesian hub identifications for XOR epistasis and our conservative pruning and identification strategy. Despite this, the ways in which these loci interact are mostly distinct in rats. This is explained by the many possible ways in which epistasis can theoretically occur which Li and Reich illustrate in their enumeration of possible full penetrance models [42]. For two-way interactions, Cartesian encodings model epistasis multiplicatively where the slope of one MLG is zero while another is double that of a third intermediate MLG (File S2). As we have shown, these interactions are indeed possible and detectable. However, genetic systems can be complex, leading to an array of possible interaction mechanisms, like in XOR (File S2). Li and Reich only highlight full penetrance models that serve as adequate models for detecting non-linear or non-additive interactions. It may not be possible for a Cartesian interaction term to describe the many types of epistatic relationships possible among loci. Our results in mice even suggest that the Cartesian encoding may not be suitable to detect epistasis in that system. While this may also be true for the XOR encoding, our results illustrate that the XOR encoding may describe more epistatic events than Cartesian in two systems.

These results suggest that multiple models of epistasis should be considered when investigating interactions among loci. Despite XOR being an extreme model of epistasis we provide statistical evidence for epistatic interactions in two natural, model systems that are detected under a full penetrance XOR model. This statistical evidence may justify experimental work to validate that biological interactions following an XOR-like model can evolve and be maintained in natural systems. The application of other penetrance functions to determine the degree of overlap between interaction encodings using our methodology may also warrant further investigation. It is likely that the XOR model, as with the Cartesian model, can only detect a portion of all epistatic events occurring across a genome.

### Genes near XOR-Unique Loci are Associated with Metabolism in rats

In rats functional annotation of gene models 1Mb upstream and downstream of the 16 loci unique to the Cartesian encoding are not enriched for any GO terms and only one KEGG pathway (Ribosome). However, gene models near the 25 loci unique to XOR yield enrichment results primarily associated with metabolism and catalysis. Of these enrichments, a KEGG pathway of primary interest is the glucagon signaling pathway as glucagon-like-protein-1 receptor agonists (GLP-1 RAs) are emerging as important medications to lower plasma glucose and induce weight loss in Murine models and humans [66, 67]. SNPs near the genes that enriched this pathway were not implicated in the associated GWAS and could present possible targets for further study with additional experimentation. When we investigate enrichment in the 75 epistatic loci shared between Cartesian and XOR encodings and the 18 putative QTL from the GWAS, immunity-related processes make up most enriched GO terms and pathways. In mice, XOR-specific GO term enrichments are mostly centered around immunity, with some notable metabolic exceptions. Since there were no Cartesian-specific epistatic loci in mice, we could not investigate Cartesian-specific enrichment. However, one of the two loci shared between encodings, chr12.7079769, is in close proximity to the *ApoB* gene. *ApoB* is an apolipoprotein and thus a structural component involved with the formation and synthesis of low density lipoproteins [68]. It has been shown that heterozygous mice for a knockout in *ApoB* were protected against diet-induced hypercholesterolemia after being fed a diet rich in fat and cholesterol [68]. Further, in a more recent study, disruption of *ApoB* led to high incidences of non-alcoholic fatty liver disease [69].

Variants in this gene are likely linked to changes in cholesterol/lipid metabolism and BMI. In rats, the XOR encoding uncovers an enrichment signal for metabolism that would have been missed if only the Cartesian encoding was applied. In contrast, in mice, strong metabolic signal is captured using the Cartesian encoding. However, numerous examples of epistasis and notable metabolic-related enrichments are also detected with XOR. It is important to note that because so many XOR-specific epistatic loci are discovered in mice, enrichment becomes more challenging as many gene models are considered, potentially weakening enrichment signals. It is difficult to extrapolate if a similar enrichment of immunity across shared loci, as we observe in rats, would have occurred if more epistasis was detected using the Cartesian encoding in mice. More experiments across breeding designs, phenotypes, and species are needed to better understand if certain interaction encodings are associated with specific biological enrichments.

Although links between immunity and obesity-related phenotypes have been well-documented [70–75], immunity-related enrichments and associations are commonly reported across diverse taxa and phenotypes [73, 76–79]. This is largely due to the inherent complexity of gene networks underlying immune systems across animal taxa [80], not excluding invertebrates [76, 78, 79]. Additionally, and perhaps even more importantly, genes related to immunity are likely core to general stress responses in all forms of life [71, 73, 77]. Since most phenotypes of interest to biologists and clinicians center around extreme perturbations from homeostasis (stressful conditions), it may be common to see the over-representation of immunity-related GO terms in functional annotations across biology and medicine. Our results may suggest that, in rats, shared epistatic loci between encodings capture the dynamic and feedback-regulated nature of immunity observed in biological systems. Yet, in a closely-related species (*M. musculus*), XOR-specific epistatic loci uncover a mostly immunity-related signal, again with notable metabolic exceptions. An important consideration is that the mice and rats used in the associated experiments, despite being closely-related biological systems, were reared in different environments, were exposed to unique stressors, ate different diets, and came from different pedigrees. One plausible explanation for the difference in enrichment profiles is the role of gut microbiota in the association between immunity and obesity. There is strong evidence in the literature of the link between the gut microbiome and immunity in affecting obesity-related phenotypes [81–84]. Furthermore there have been notable differences identified between rats and mice concerning the role of the microbiome in immunity and obesity [85, 86]. It may be possible that our distinct enrichment profiles between systems may be highlighting species-level differences in how obesity is related to immunity. Alternatively, we could also be capturing signals associated with differences between methodologies and/or environments. Additional experiments are required to elucidate the differences in how epistatic interactions underlie obesity in these species.

In rats, the XOR model identifies epistatic loci enriched for more biologically relevant functions and processes. The XOR model yields enrichment more closely associated with metabolism and BMI. Full penetrance XOR logic may not be biologically plausible due to genetic constraints. However, a possible scenario in which it may occur is when the presence of one activator (A), encoded by gene one, while in the presence of another activator (B), encoded by gene two, results in the transcription of gene three. However, in the presence of both activators, gene three is not expressed.

This could be due to activators A and B binding to one another when they co-occur, inhibiting their DNA binding motifs, or because both activators collectively block other transcription enzymes from binding [44]. Mechanistic examples of XOR interactions in transcriptional regulation such as this may be plausible. However, examples of XOR logic in other biological processes may seem less likely. The XOR model assumes a phenotypic score in one extreme when only one locus is in a heterozygous state but the other extreme if both loci are heterozygous (File S2). In terms of BMI, where higher phenotypic scores are deleterious in a static environment where food is abundant, this would translate to heterozygote disadvantages in one locus when the other locus is in a homozygous state. However, if both loci are in a heterozygous state, then the MLG is associated with lower BMI, leading to a heterozygote advantage. Heterotic relationships are commonly described in natural systems, primarily in crop plants [87] but also in humans as observed in the genetic mechanisms underlying sickle-cell anemia and malaria resistance [88]. Yet, the variable phenotypic diversity of sickle cell morphology and/or malaria resistance violates Mendelian expectations [11]. It is possible that one explanation for these abnormalities are epistatic interactions between loci, including relationships described by penetrance functions other than Cartesian. Exploring these phenotypes with multiple interaction encodings may assist in explaining deviations from Mendelian expectations in malaria and other phenotypes.

### Conclusions and Future Work

Regardless of the underlying mechanisms, our results show that in our rat data, some epistatic loci are located near genes that are enriched for metabolic functions and interact in a manner more detectable by the XOR model. In our mouse samples, we illustrate that many epistatic events underlying BMI are only detectable using an XOR model. It is important to keep in mind that we are applying a full penetrance model to a continuous phenotype (BMI). While the conventional XOR model assumes a binary, discrete outcome of presence or absence of a certain phenotypic state (commonly a disease phenotype), we have used the XOR encoding with a continuous phenotype as a example alternative model for the Cartesian encoding and were able to detect many pairwise and three-way interactions between SNPs. Our two-way and three-way results in rats suggest that epistatic loci interact in different ways depending on the interaction encoding used.

Distinct interaction mechanisms may exist for different biological networks involving shared loci. Taken together, our study has been illustrative in showing that epistatic interactions in biological systems are likely far more complex and ubiquitous than previously thought. This necessitates the consideration of different models of interaction for investigating epistasis. The algorithms we have given provide tools for collecting statistical evidence for epistasis using different models of interaction. The matrix-based permutation testing algorithm we have presented can give further statistical evidence and can also be simplified and applied in GWAS or eQTL studies. In the future, we will apply our methodology to diverse taxa and phenotypes to investigate the complexity of epistasis and warrant the development of validation studies to describe biological interactions. We hope to further elucidate complex traits, including many human diseases, by uncovering genetic relationships that have thus far been elusive to standard analyses. Extending the algorithms given for logistic regression for case/control studies and generalized linear models would be very useful and important especially for handling population structure directly. Li and Reich presented many different models for epistasis and corresponding penetrance functions [42]. A natural extension to this work would be to use more penetrance functions for encoding the interactions. This can already be done with the algorithms given with the only change being how the interaction terms are encoded according to the penetrance function. The work for this software extension is already under development.

## Supporting information

GWAS and epistastic subsets raw data

Reaction norms

Loci counts and enrichment

## Acknowledgements

We would like to thank Drs. Takeshi Matsui (Stanford), Sasha Levy (Stanford), and Ian Ehrenreich (University of Southern California, Dornsife) for providing R scripts and data to generate interaction plots in Figures 2, 6, and 8. We would also like to thank Dr Karl Broman (University of Wisconsin, Madison) for his suggestions in handling the relatedness structure in the rat data set.

## Data Availability

Rat phenotype data and GWAS summary statistics are available at https://library.ucsd.edu/dc/object/bb83725195. Rat genotype data are available at: https://library.ucsd.edu/dc/object/bb15123938. Mouse genotype and phenotype data are available via the “BGLR” package in R.

## Footnotes

1 https://github.com/EpistasisLab/epistasis_detection

