## Supplementary material for "Computational framework for statistical epistasis supports XOR penetrance function in a living system": Reaction norms

**File S2**

I. Additive, non-epistatic two-way model.

a. A completely additive, non-epistatic model between two loci. Numbers in parenthesis are the genotype encodings used in the raw data.

|  |  | **SNP2** | | |
| --- | --- | --- | --- | --- |
|  |  | **BB (0)** | **Bb (1)** | **bb (2)** |
| **SNP1** | **AA (0)** | 0 | 1 | 2 |
|  | **Aa (1)** | 1 | 2 | 3 |
|  | **aa (2)** | 2 | 3 | 4 |

b. Theoretical reactions norms for an additive, non-epistatic model between two loci. Colored lines represent the reaction norms for SNP1 genotypes (see legend). SNP2 genotypes are on the x-axis. Hypothetical phenotypic scores are on the y-axis.

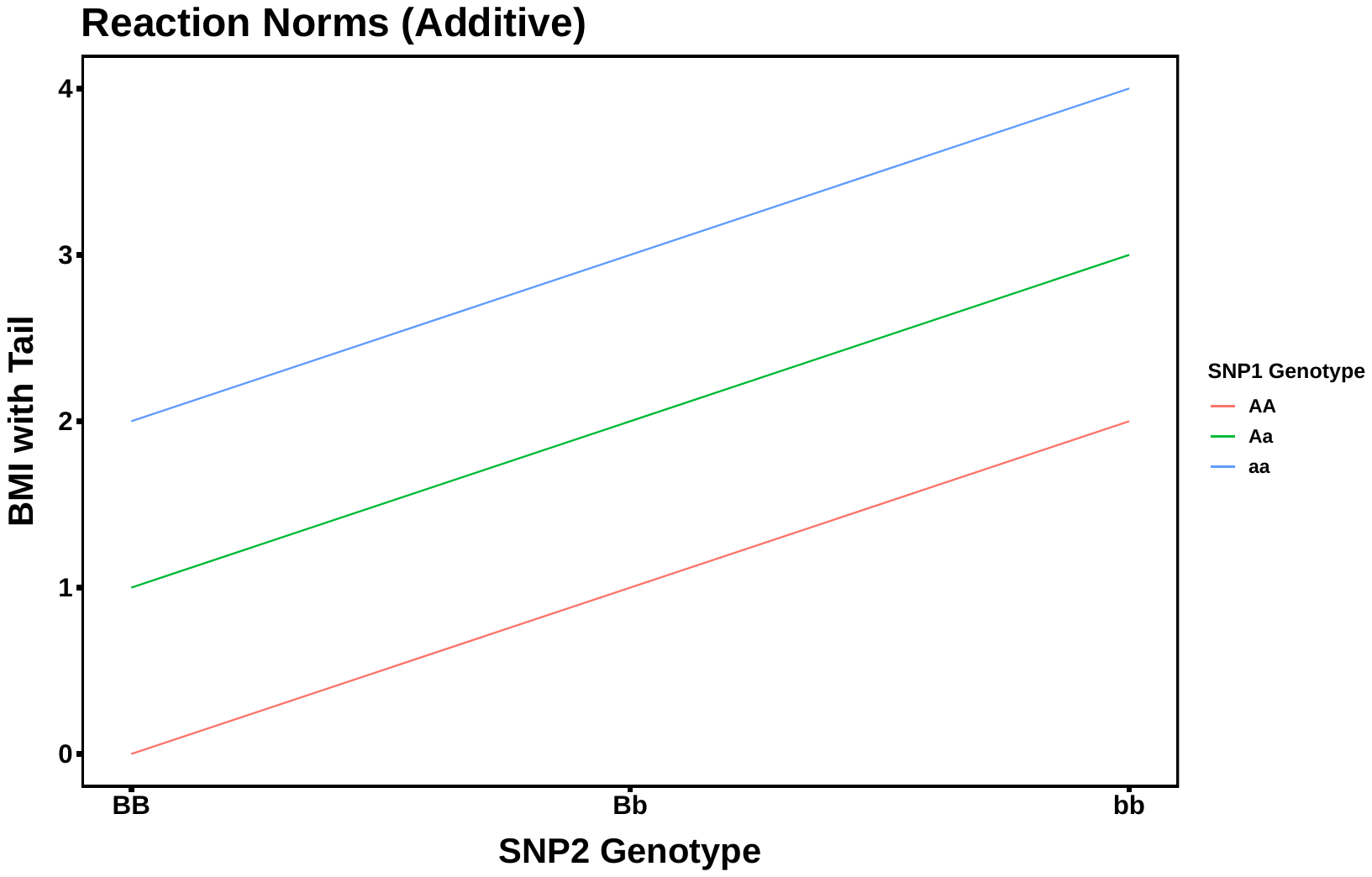

c. Example reactions norms for a near additive, non-epistatic interaction between two loci found in the rat dataset. Colored lines represent the reaction norms for SNP1 (chr1:202370696_C) genotypes (see legend). SNP2 (chr3:109942006_C) genotypes are on the x-axis. Actual phenotypic scores are on the y-axis. Note: sign of slopes are reversed compared to theoretical example.

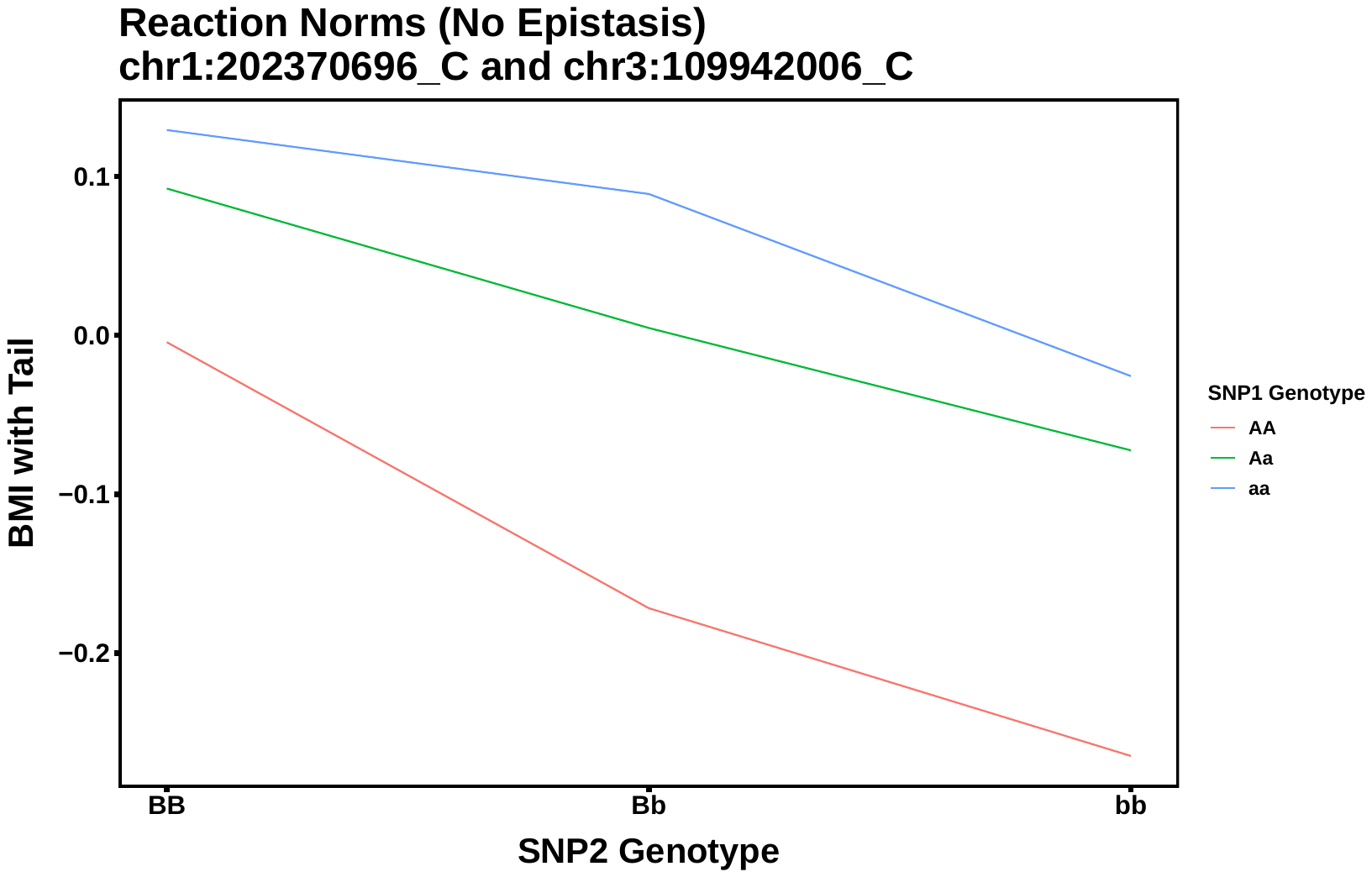

II. Cartesian two-way model of epistasis.

a. The Cartesian model of epistasis. Numbers in parenthesis are the genotype encodings used in the raw data.

|  |  | **SNP2** | | |
| --- | --- | --- | --- | --- |
|  |  | **BB (0)** | **Bb (1)** | **bb (2)** |
| **SNP1** | **AA (0)** | 0 | 0 | 0 |
|  | **Aa (1)** | 0 | 1 | 2 |
|  | **aa (2)** | 0 | 2 | 4 |

b. Theoretical reaction norms of the Cartesian model between two loci. Red and blue dashed lines are the reaction norms for the homozygous genotypes for SNP1. Green dashed line represents the reaction norm for the heterozygous genotype for SNP1. SNP2 genotypes are on the x-axis. Hypothetical phenotypic scores are on the y-axis.

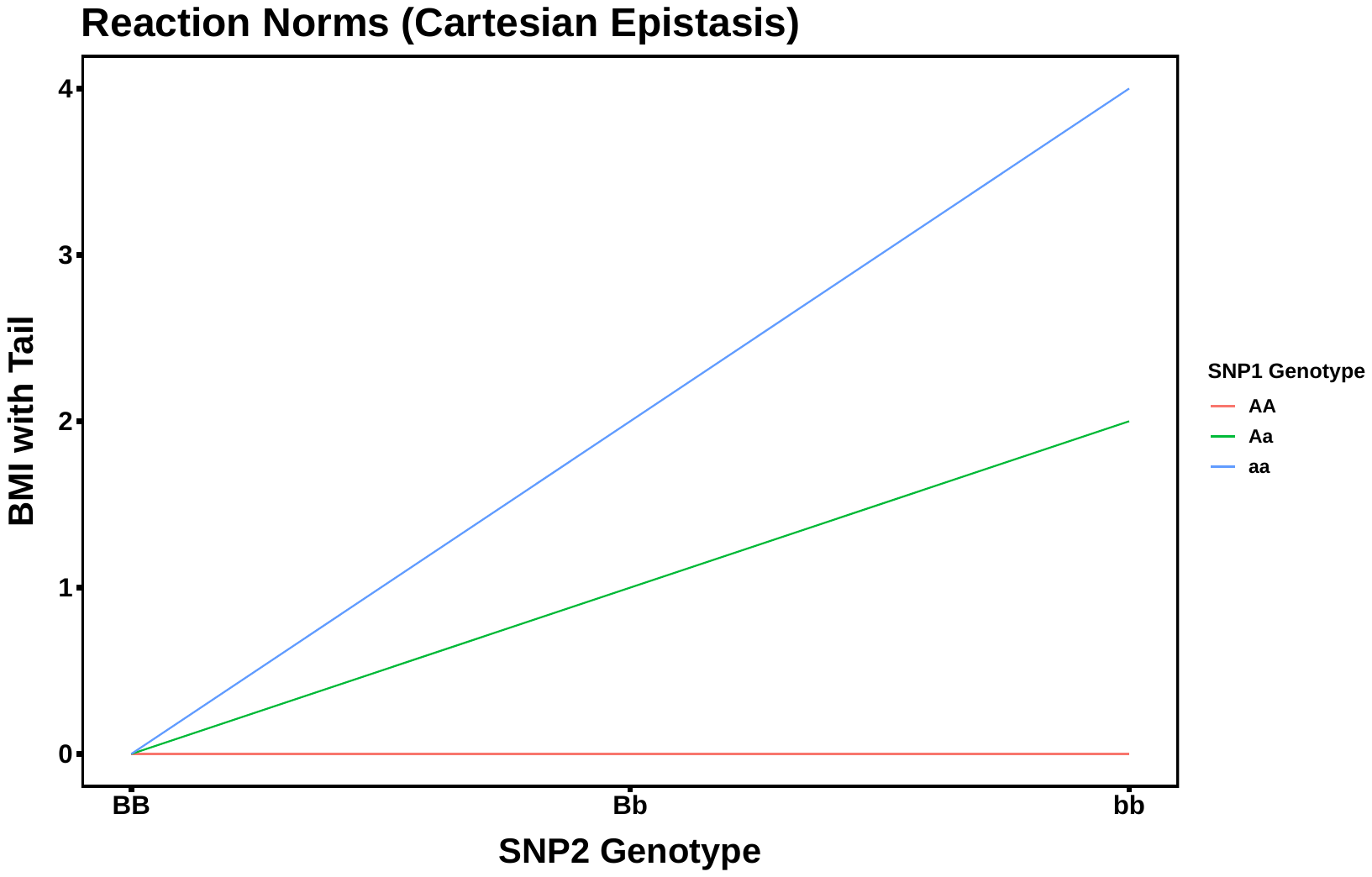

c. Example reactions norms for a near Cartesian interaction between two loci found in the rat dataset. Colored lines represent the reaction norms for SNP1 (chr10:84091208_T) genotypes (see legend). SNP2 (chr12:29243508_C) genotypes are on the x-axis. Actual phenotypic scores are on the y-axis.

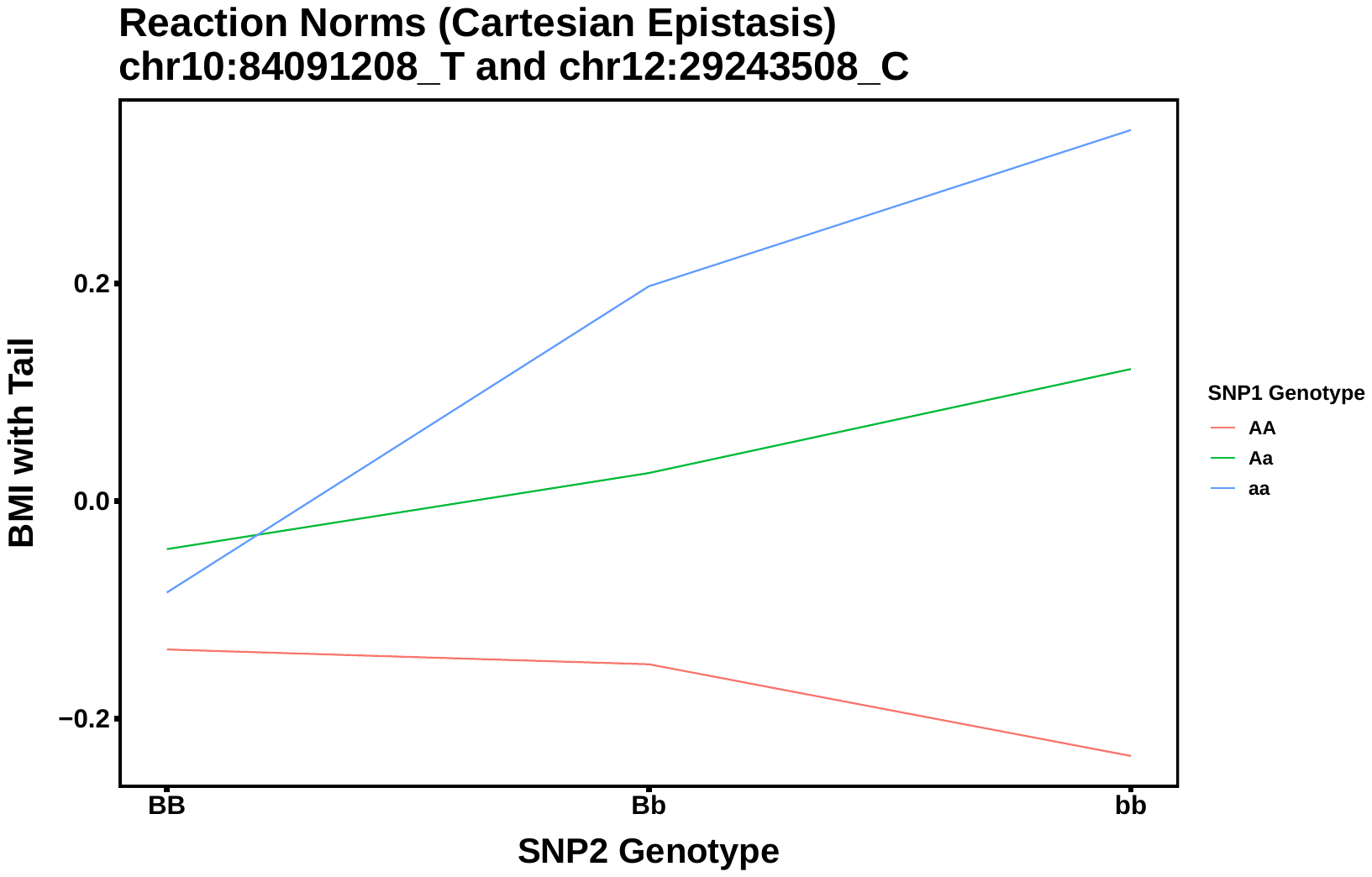

d. Example reactions norms for a near Cartesian interaction between two loci found in the mouse dataset. Colored lines represent the reaction norms for SNP1 (chr3:44652325_G) genotypes (see legend). SNP2 (chr12:7079769_C) genotypes are on the x-axis. Actual phenotypic scores are on the y-axis.

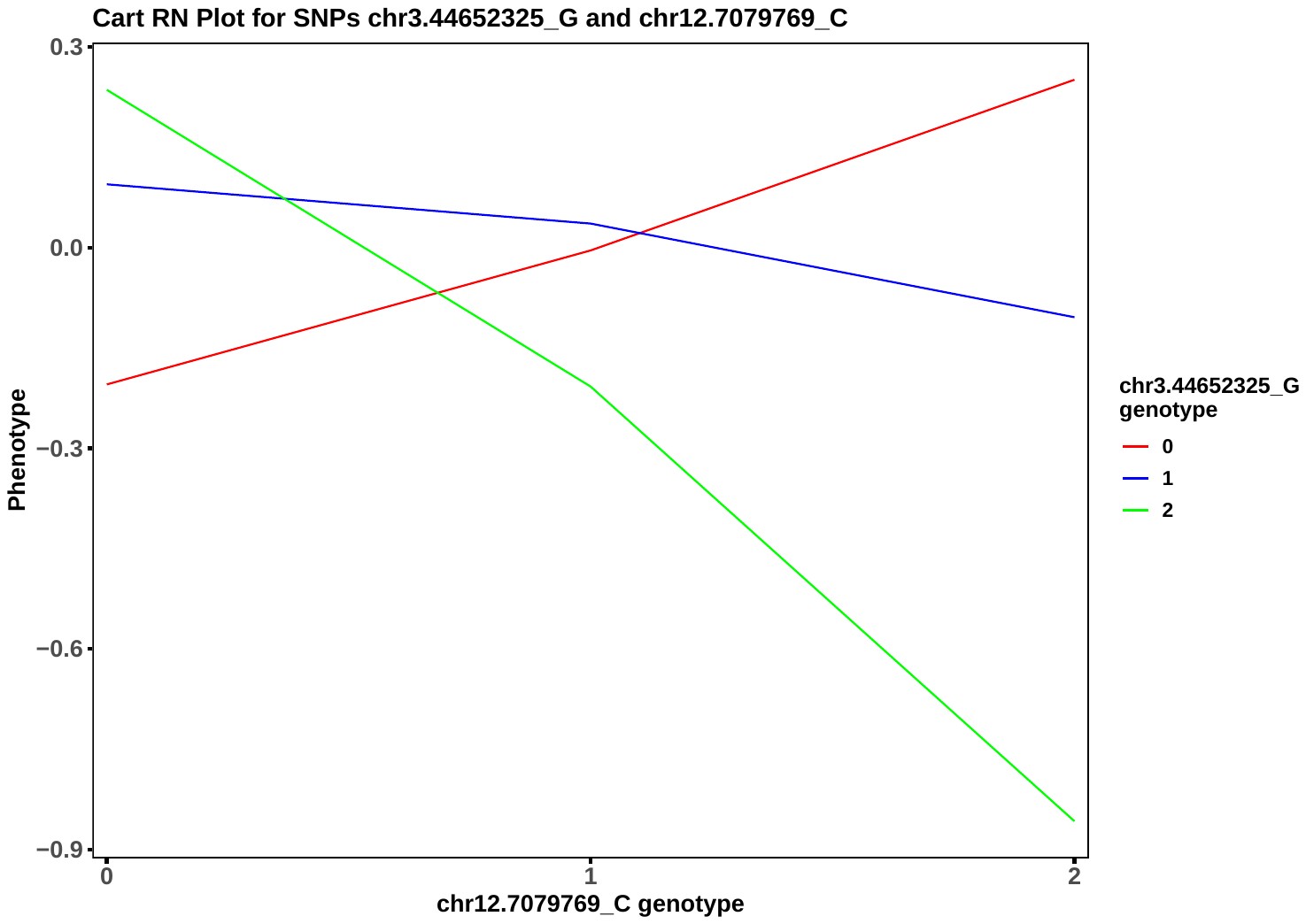

III. XOR full penetrance two-way model of epistasis.

a. The XOR full penetrance model of epistasis (1). Marginal penetrance values represent phenotypic scores (or disease states) that are equal among single genotypes. Numbers in parenthesis are the genotype encodings used in the raw data.

|  | | **SNP2** | | | |
| --- | --- | --- | --- | --- | --- |
|  |  | **BB (0)** | **Bb (1)** | **bb (2)** | **Marginal Penetrance** |
| **SNP1** | **AA (0)** | 0 | 1 | 0 | 0.5 |
|  | **Aa (1)** | 1 | 0 | 1 | 0.5 |
|  | **aa (2)** | 0 | 1 | 0 | 0.5 |
|  | **Marginal Penetrance** | 0.5 | 0.5 | 0.5 |  |

b. Theoretical reaction norms of the XOR model between two loci. Red and blue dashed lines are the reaction norms for the homozygous genotypes for SNP1. Green dashed line represents the reaction norm for the heterozygous genotype for SNP1. SNP2 genotypes are on the x-axis. Hypothetical phenotypic scores are on the y-axis.

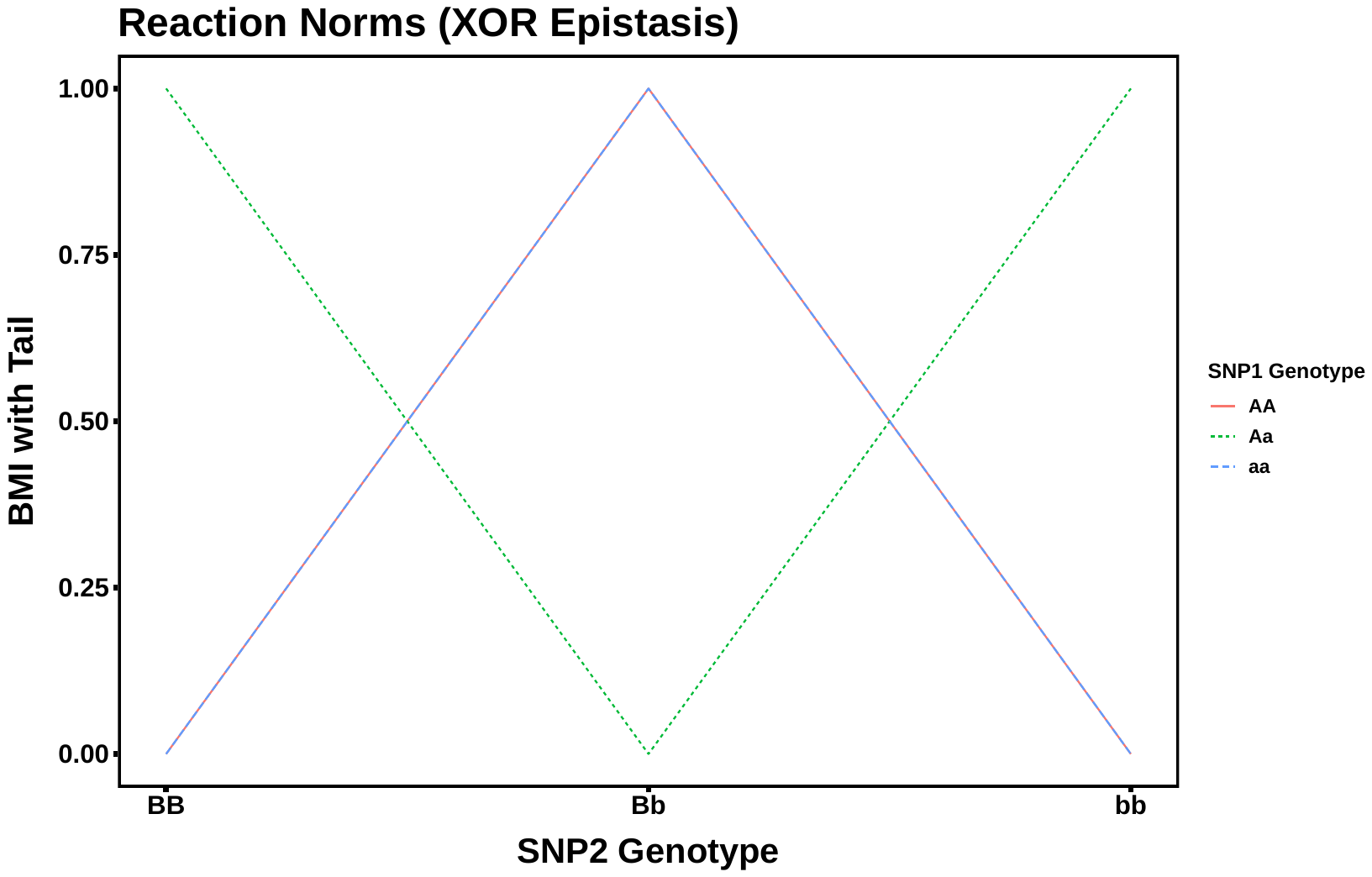

c. Example reactions norms for an XOR interaction between two loci found in rat dataset. Colored lines represent the reaction norms for SNP1 (chr2:48769279_C) genotypes (see legend). SNP2 (chr9:53532432_C) genotypes are on the x-axis; phenotypic scores on y-axis.

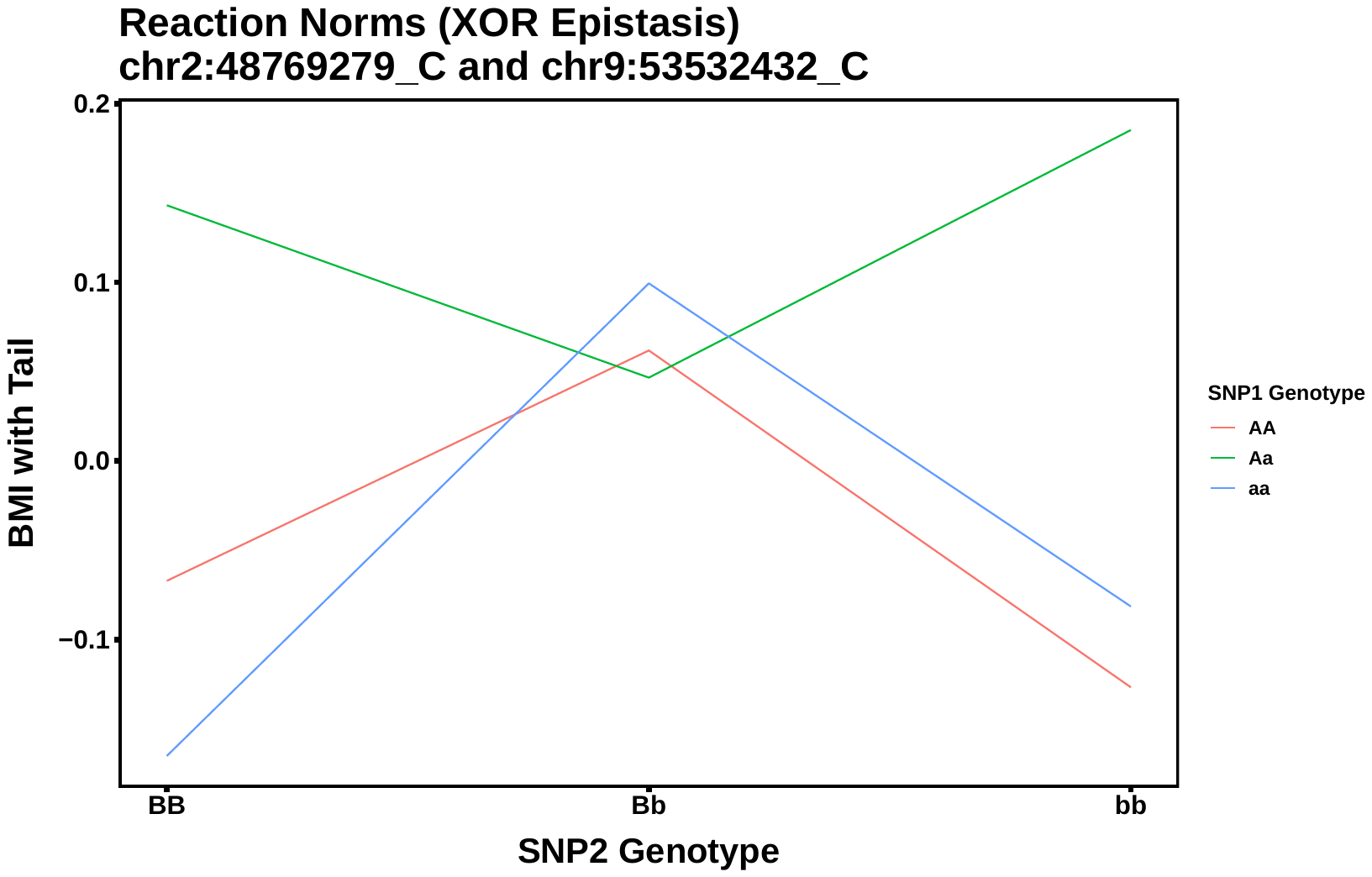

c. Example reactions norms for an XOR interaction between two loci found in mouse dataset. Colored lines represent the reaction norms for SNP1 (chr6:43058217_A) genotypes (see legend). SNP2 (chr13:27083871_A) genotypes are on the x-axis; phenotypic scores on y-axis.

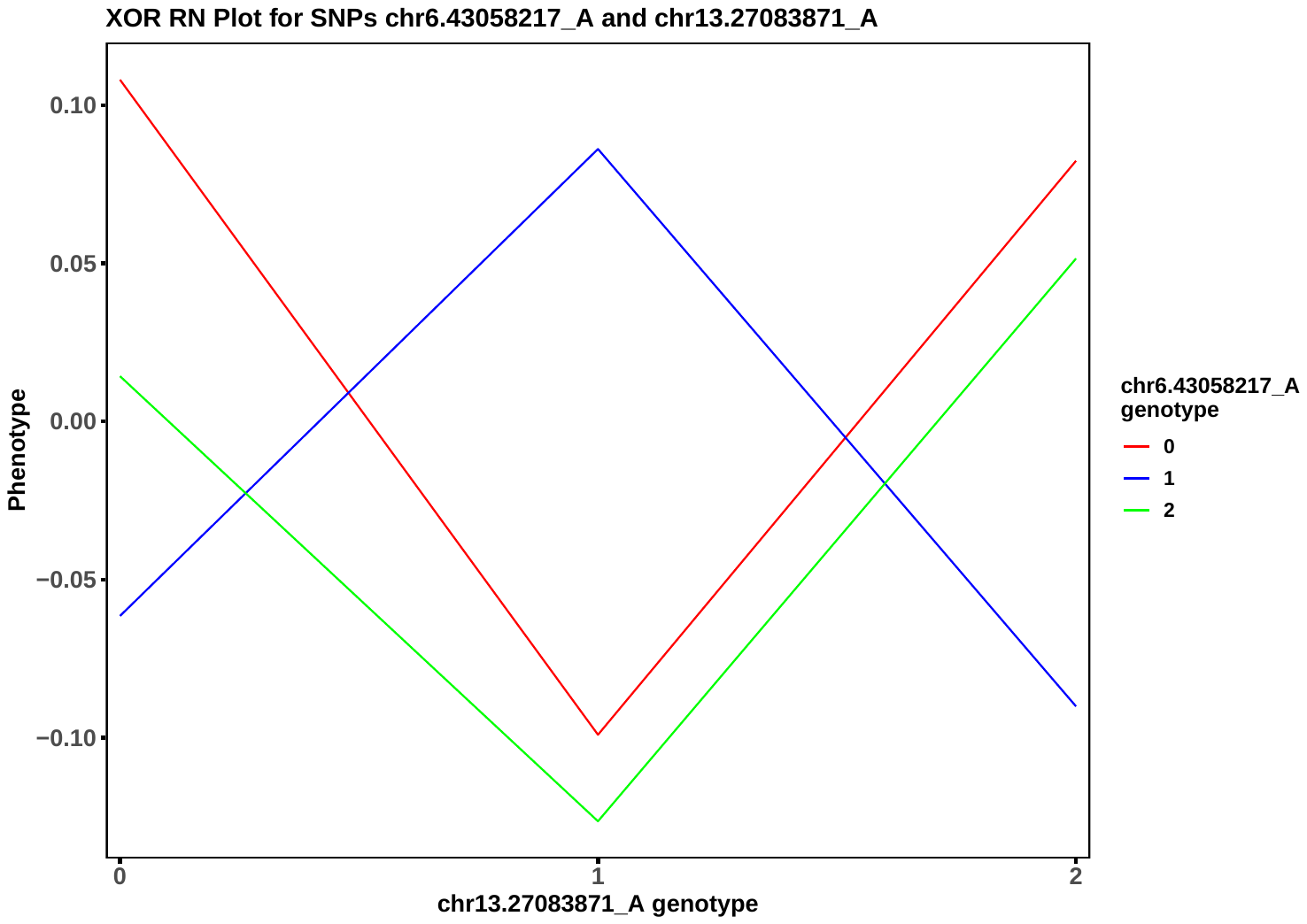

IV. Three-way Epistatic models.

a. Additive three-way, non-epistatic model. Numbers in parenthesis are the genotype encodings used in the raw data.

|  | | **SNP3** | | | | | | | | |
| --- | --- | --- | --- | --- | --- | --- | --- | --- | --- | --- |
|  |  | **CC (0)** | | | **Cc (1)** | | | **cc (2)** | | |
|  |  | **SNP2** | | | **SNP2** | | | **SNP2** | | |
|  |  | **BB (0)** | **Bb (1)** | **bb (2)** | **BB (0)** | **Bb (1)** | **bb (2)** | **BB (0)** | **Bb (1)** | **bb (2)** |
| **SNP1** | **AA (0)** | 0 | 1 | 2 | 1 | 2 | 3 | 2 | 3 | 4 |
|  | **Aa (1)** | 1 | 2 | 3 | 2 | 3 | 4 | 3 | 4 | 5 |
|  | **aa (2)** | 2 | 3 | 4 | 3 | 4 | 5 | 4 | 5 | 6 |

b. Cartesian three-way, epistatic model. Numbers in parenthesis are the genotype encodings used in the raw data.

|  | | **SNP3** | | | | | | | | |
| --- | --- | --- | --- | --- | --- | --- | --- | --- | --- | --- |
|  |  | **CC (0)** | | | **Cc (1)** | | | **cc (2)** | | |
|  |  | **SNP2** | | | **SNP2** | | | **SNP2** | | |
|  |  | **BB (0)** | **Bb (1)** | **bb (2)** | **BB (0)** | **Bb (1)** | **bb (2)** | **BB (0)** | **Bb (1)** | **bb (2)** |
| **SNP1** | **AA (0)** | 0 | 0 | 0 | 0 | 0 | 0 | 0 | 0 | 0 |
|  | **Aa (1)** | 0 | 1 | 2 | 0 | 1 | 2 | 0 | 2 | 4 |
|  | **aa (2)** | 0 | 2 | 4 | 0 | 2 | 4 | 0 | 4 | 8 |

c. XOR full penetrance three-way, epistatic model. Numbers in parenthesis are the genotype encodings used in the raw data.

|  | | **SNP3** | | | | | | | | |
| --- | --- | --- | --- | --- | --- | --- | --- | --- | --- | --- |
|  |  | **CC (0)** | | | **Cc (1)** | | | **cc (2)** | | |
|  |  | **SNP2** | | | **SNP2** | | | **SNP2** | | |
|  |  | **BB (0)** | **Bb (1)** | **bb (2)** | **BB (0)** | **Bb (1)** | **bb (2)** | **BB (0)** | **Bb (1)** | **bb (2)** |
| **SNP1** | **AA (0)** | 0 | 1 | 0 | 1 | 0 | 1 | 0 | 1 | 0 |
|  | **Aa (1)** | 1 | 0 | 1 | 0 | 1 | 0 | 1 | 0 | 1 |
|  | **aa (2)** | 0 | 1 | 0 | 1 | 0 | 1 | 0 | 1 | 0 |

V. Statistics for correlation tests (cor.test in R) in rats between GWAS metrics (-log10*P* or absolute value of beta coefficient) and number of significant epistatic interactions detected for each epistatic locus or QTLs only under both Cartesian and XOR encodings for both two-way and three-way experiments. “**r**” in the table is the Pearson’s product-moment correlation.

| ***GWAS Metric*** | ***Test Statistic*** | ***2w-Cartesian*** | ***2w-Cartesian (QTL only)*** | ***2w-XOR*** | ***2w-XOR (QTL only)*** | ***3w-Cartesian (QTL only)*** | ***3w-XOR (QTL only)*** |
| --- | --- | --- | --- | --- | --- | --- | --- |
| ***-log_10_P*** | **t** | 1.242 | 1.782 | 2.586 | 3.583 | 0.542 | 1.893 |
|  | **df** | 89 | 14 | 98 | 16 | 16 | 16 |
|  | ***p*-value** | 0.218 | 0.0965 | **0.0112** | **0.00249** | 0.595 | 0.0766 |
|  | **95% CI** | (-0.0775, 0.328) | (-0.0836, 0.763) | **(0.0593, 0.428)** | **(0.291, 0.865)** | (-0.355, 0.566) | (-0.0489, 0.746) |
|  | **r** | 0.130 | 0.430 | **0.253** | **0.667** | 0.134 | 0.428 |
| ***abs(beta)*** | **t** | -0.187 | 0.795 | 0.915 | 1.217 | -0.194 | -0.254 |
|  | **df** | 89 | 14 | 98 | 16 | 16 | 16 |
|  | ***p*-value** | 0.852 | 0.440 | 0.363 | 0.241 | 0.849 | 0.803 |
|  | **95% CI** | (-0.225, 0.187) | (-0.321, 0.638) | (-0.106, 0.283) | (-0.203, 0.667) | (-0.504, 0.428) | (-0.515, 0.416) |
|  | **r** | -0.0198 | 0.208 | 0.0920 | 0.291 | -0.0483 | -0.632 |

VI. Dotplots for rats for enriched GO Terms (Biological Process and Molecular Function) for the gene models derived from the 18 putative QTL from the original rat GWAS [2,3].

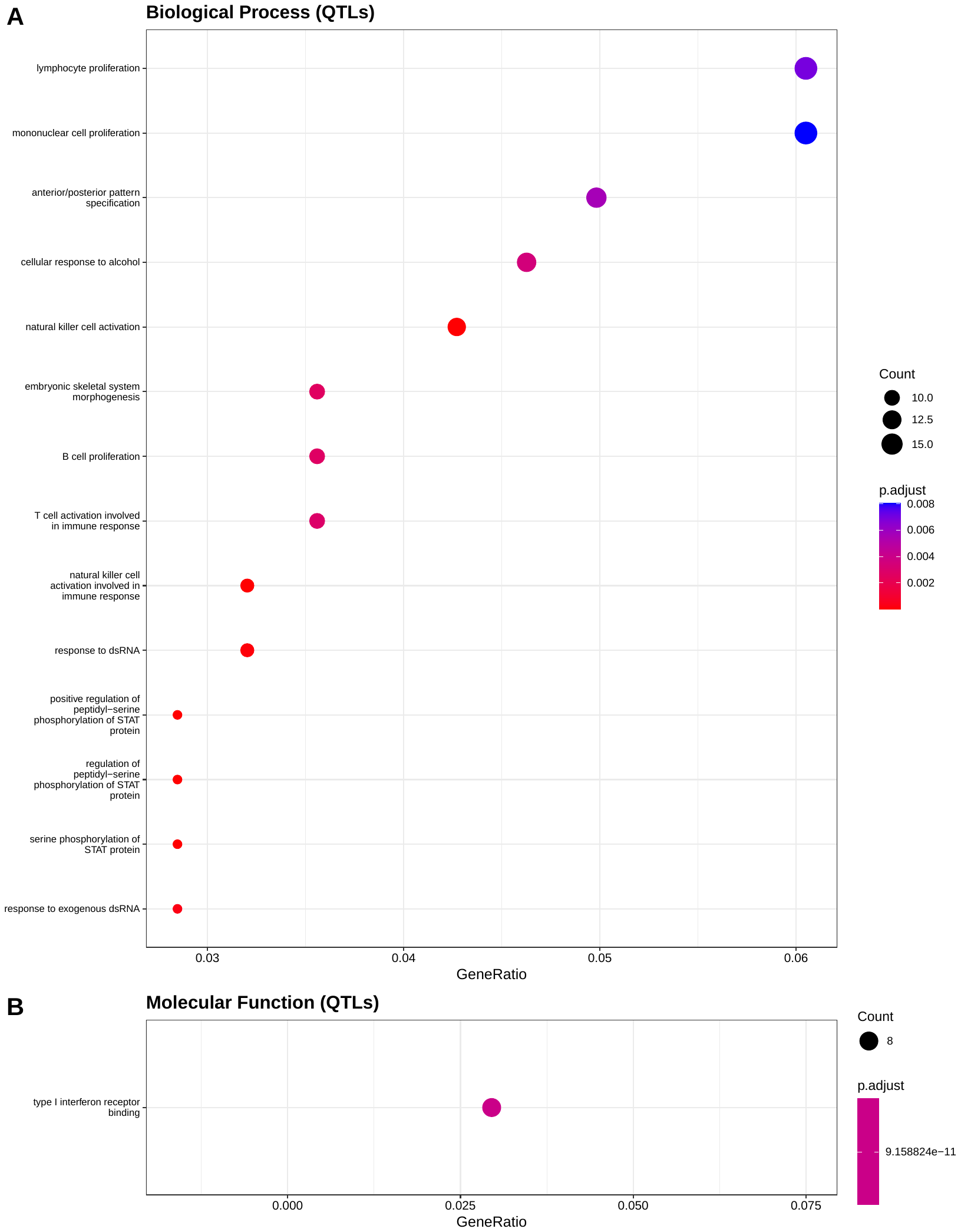

VII. Dotplots for mice for enriched GO Terms (Cellular Component) for the gene models derived from the 3 putative QTL from the original mouse GWAS [4].

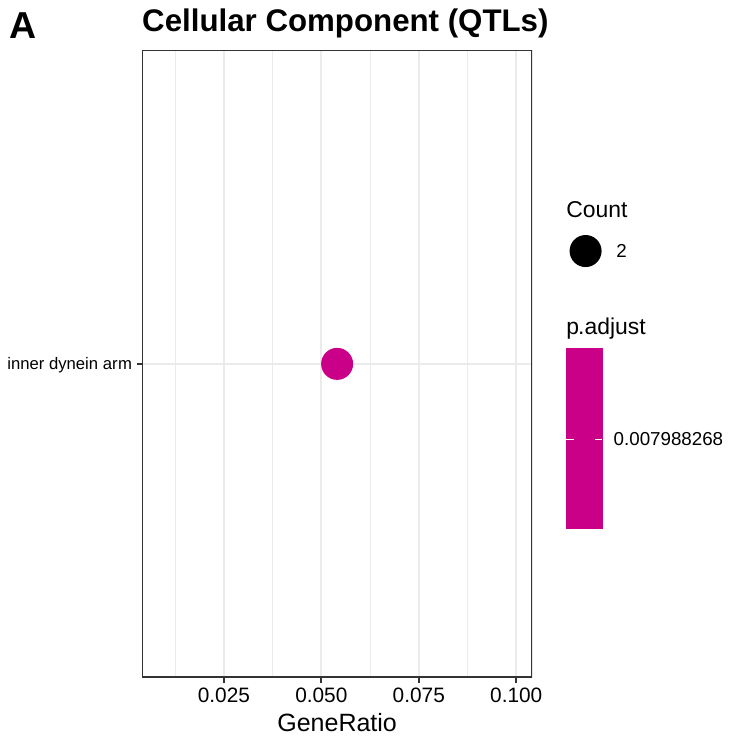
